# A human and mouse subpopulation of senescent β-cells induces pathologic dysfunction through targetable paracrine signaling

**DOI:** 10.1101/2025.05.07.648438

**Authors:** Kanako Iwasaki, Priscila Carapeto, Cristian Abarca, Francesko Hela, Stephanie Sanjines, Sebastian Pena, Sandra Le, Hui Pan, Christopher Cahill, Ayush Midha, Juliana Alcoforado Diniz, Dylan Baker, Sergii Domanskyi, Sara Espinoza, Alejandro Pena, Francisco G. Cigarroa, Jillian L. Woolworth, Jeffrey H. Chuang, Vesna D Garovic, James L. Kirkland, Tamar Tchkonia, Nicolas Musi, George A. Kuchel, Paul Robson, Cristina Aguayo-Mazzucato

**Author notes:** These authors contributed equally to this work (Kanako Iwasaki, Priscila Carapeto).

## Abstract

Cellular senescence is a stress response mechanism marked by irreversible growth arrest, upregulation of antiapoptotic pathways, loss of cellular function, and remodelling of the cellular secretory profile. In both humans and mice, pancreatic β-cells undergo senescence with age and insulin resistance. Targeted removal of senescent cells in mouse models of diabetes improves glucose homeostasis, demonstrating the role β-cell senescence in diabetes progression. In contrast, β-cell senescence also promotes immune surveillance, promoting β-cell survival and function. Thus, a better understanding of senescent cells’ phenotypic and functional heterogeneity is needed to develop effective therapeutic strategies.

Herein, we show that subpopulations of senescent β-cells in mice and humans, which were identified through the expression of *Cdkn1a* (encoding *p21^Cip1^*) and *Cdkn2a* (encoding *p16^Ink4a^*) by single-cell RNA sequencing (scRNA-seq), flow cytometry, spatial transcriptomics, and spatial proteomics, exhibit distinct transcriptional and functional identities. The predominant senescent β-cell subpopulation expressed *Cdkn1a* and was characterized by a lack of glucose responsiveness, high basal insulin secretion, and transcription of canonical SASP factors. The SASP of *Cdkn1a*-expressing β-cells had non-cell autonomous effects on neighbouring cells. A subset of four SASP factors from *Cdkn1a^+^* cells was sufficient to induce secondary senescence and β-cell dysfunction *in vitro*. JAK inhibitors (JAK1/2 and JAK1/3) counteracted secondary senescence induction and restored β-cell function in high-fat diet-fed mice and human islets from donors with or without type 2 diabetes.

**GRAPHICAL ABSTRACT:** 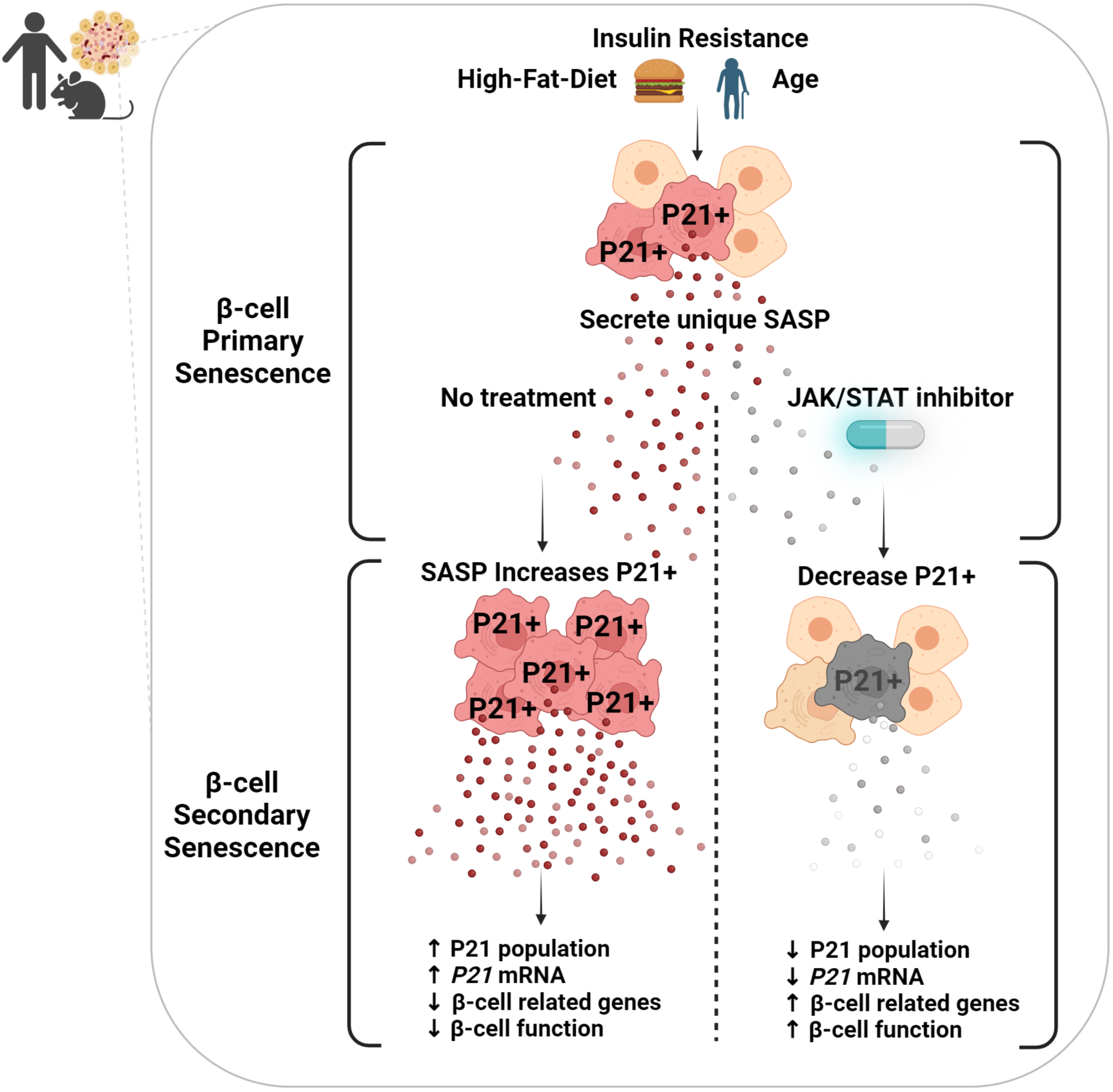

## INTRODUCTION

As cells age or undergo stress, they may enter a state known as cellular senescence, characterized by a halt in cellular division, increased cellular dysfunction, and upregulation of antiapoptotic pathways while maintaining an active metabolic profile characterized by secretion of senescence-associated secretory phenotype (SASP) factors. The non-cell autonomous effects of the SASP include the induction of secondary senescence in surrounding cells [1], thereby amplifying the detrimental effects of senescence. Pharmacological and nonpharmacological interventions have been developed to reduce the load and deleterious effects of senescent cells. Senolytics promote death and clearance of senescent cells, while senomorphics target different SASP-regulating pathways to reduce the release of SASP factors and their paracrine and systemic effects.

Insulin resistance and type 2 diabetes (T2D) are associated with increased senescence markers in mouse and human pancreatic β-cells [2, 3]. Our prior studies revealed that senescent cells are dysfunctional (characterized by high basal insulin secretion and lack of glucose responsiveness) and secrete a unique senescence-associated secretory phenotype (SASP) enriched in inflammatory signals and extracellular matrix remodeling factors [3–6]. Senolysis improved insulin sensitivity [7], glucose homeostasis, and β-cell identity and function while decreasing the transcription of SASP factors in mouse models [3]. Nonetheless, the effect of SASP factors on secondary senescence remains to be determined. A complementary approach to targeting β-cell senescence is through its SASP. The Janus kinase/signal transducer and activator of transcription (JAK/STAT) pathway is one of the known regulators of cytokine production [8, 9], and its inhibition has been shown to downregulate SASP secretion in adipose tissue [10].

There are also reports of beneficial effects of β-cell senescence, such as inducing immune surveillance and promoting cell survival, differentiation, and insulin secretion [11, 12]. These contradictory results could be reconciled by a proposed heterogeneity of senescent β-cells, involving the heterogeneous distribution of senescence markers within pancreatic islets [2]. However, heterogeneity of these cells in terms of their identity, function, and secondary effects on other surrounding cells remains mostly unexplored. Efforts to define and deconvolute heterogeneity of these senescent cells will be required to develop effective and safe interventions in chronic diseases involving senescent β-cells, such as T2D.

We hypothesized that senescent cell heterogeneity is driven by differential upregulation of the cell cycle arrest genes *Cdkn1a (*encoding *p21^Cip1^)* and/or *Cdkn2a (*encoding *p16^Ink4a^),* both of which are known to mediate entry into senescence.

We identified subpopulations of senescent β-cells based on the expression of the cell cycle inhibitors *Cdkn1a* and *Cdkn2a*. The predominant senescent β-cell subpopulation in *C57BL/6* mice was *Cdkn1a*-expressing, characterized by transcription of canonical SASP factors and loss of β-cell function. In the whole human pancreas, the spatial transcriptomic and proteomic analyses demonstrated that CDKN1A/P21^CIP1+^ INS^+^ cells exhibited downregulation of key β-cell genes and transcription factors, consistent with loss of identity. Factors secreted by *Cdkn1a^+^* β-cells may have contributed to induced secondary senescence in pancreatic islets, with loss of transcriptional identity and lack of insulin secretion in response to glucose. JAK inhibitors (JAK1/2i and JAK1/3i) prevented the *Cdkn1a* SASP-induced increase in secondary senescent cells. In human islets, JAK1/2 inhibitors decreased *P21CIP1* mRNA, inhibited SASP factor release, and improved glucose responsiveness in islets from donors with and without T2D. These results reveal the non-cell autonomous effects of the β-cell SASP, which is capable of inducing secondary senescence. This process was attenuated by JAK inhibitors, which restored β-cell function and identity in both mice and humans.

## RESULTS

### p21^CIP1+^ senescent β-cells in mice are dysfunctional and lose transcriptional identity

We used previously published scRNA-Seq data [6] to analyze gene expression in pancreatic islets from adult mice following acute induction of insulin resistance with the insulin receptor antagonist S961, with a two-week recovery period **(Fig. 1A)** completely reversing hyperglycemia and hyperinsulinemia (**Suppl. Fig. 1**). Further analysis of these data revealed that β-cells, identified by the expression of *Ins2*, in 6- to 9-month-old male controls (PBS), represented 64% of the total islet population **(Fig. 1B)**. Other islet cell types were represented in the following proportions: α-11%, δ-9% PP-2%, ductal-2%, and endothelial-5% **(Suppl. Fig. 2)**. Density estimates for the expression of *p21^Cip1^* (encoded by *Cdkn1a*) or *p16^Ink4a^* (encoded by *Cdkn2a*) using a Gaussian finite mixture model identified at least two subpopulations of senescent β-cells: *Cdkn1a*^+^ and *Cdkn2a*^+^, as well as a third non-senescent of double-negative *Cdkn1a*^-^/*Cdkn2a*^-^ cells. Non-senescent (*Cdkn1a^-^/Cdkn2a^-^*) β-cells represented 75% of the total population of control islets. The remaining senescent cell subpopulations included 29% *Cdkn1a^+^* cells and 0.3% *Cdkn2a^+^* cells. These proportions align with reported values of 20-40% of *CDKN1A^+^*β-cells from adults over 60 years old [13]. The frequency of the senescent subpopulation in other islet cell types is shown in **Suppl. Table 1**.

**Figure 1.**
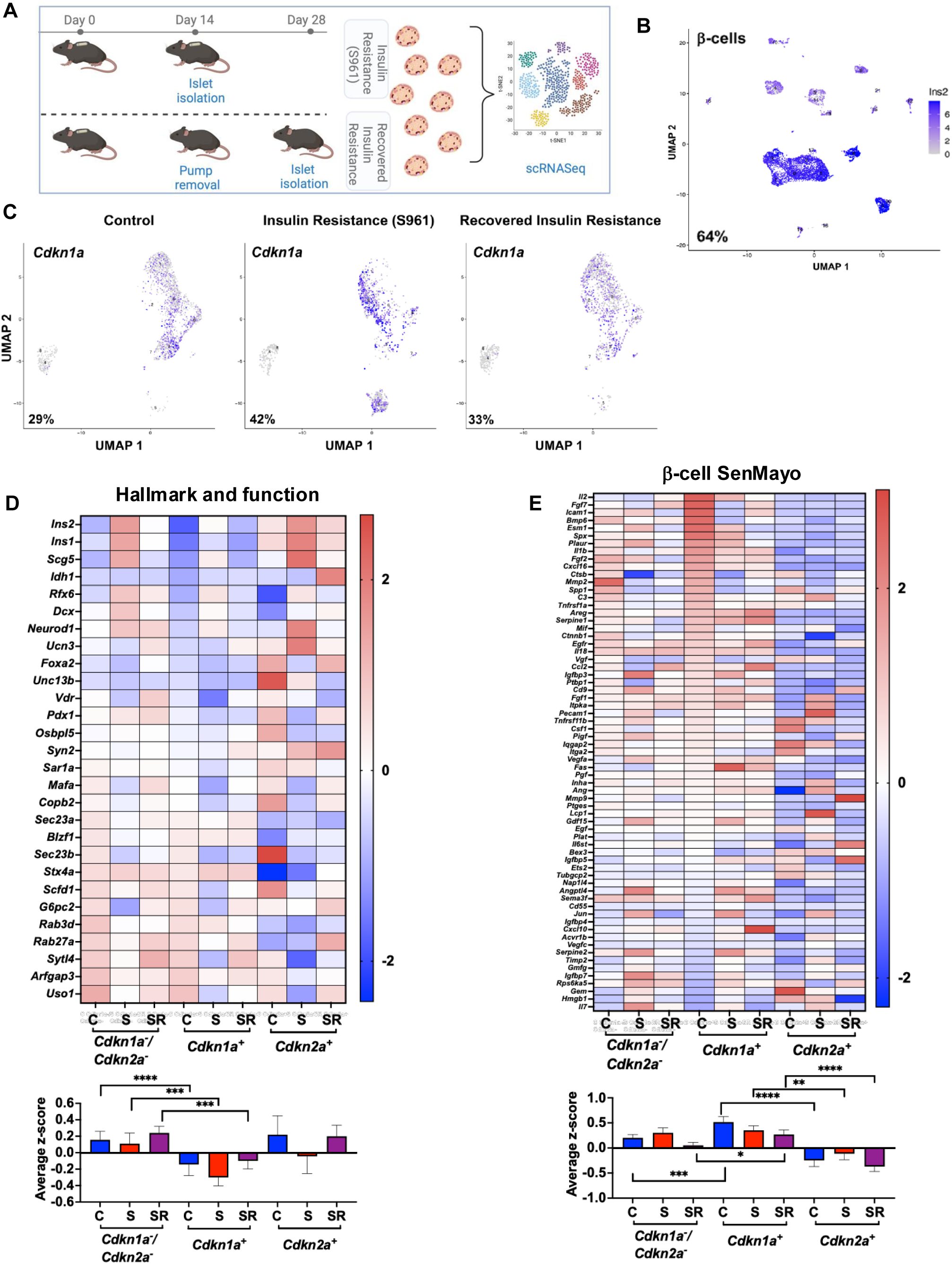
Subpopulations of senescent β-cells with different functional and SASP transcriptional profiles. **(A)** Reanalysis of scRNA-Seq data (GSE149984) of islets isolated from C57Bl/6 mice under three conditions: control (C), S961-induced insulin resistance for 2 weeks (S) and S961-induced insulin resistance followed by a 2-week recovery period (SR); **(B)** UMAP plot displaying the major islet-cell cluster of mouse β-cells based on *Ins2* expression; **(C)** UMAP showing the percentage of *Cdkn1a^+^* β-cells under the three different metabolic conditions; **(D)** Heatmap of β-cell Hallmark and Function genes. The graph shows the expression of the genes in three different cell subpopulations of non-senescent β-cells (*Cdkn1a^-^/Cdkn2a^-^*) and *Cdkn1a^+^*, *Cdkn2a^+^* under the three different metabolic conditions described. A bar graph represents the average z-score expression levels from the heatmap; Mean+/- SEM; expression levels analyzed by two-way t-tests; *p<0.01, **p<0.001 ***p<0.0001 and ****p<0.00001. Data are results from 2939 β-cells in C, 2896 β-cells from S961 conditions and 2513 cells from SR; islets were isolated from four mice per condition as previously published [1]. The midpoint of 0 represents average expression across all samples. **(E)** βSenMayo: Selected β-cell hallmark and functional genes in the same three cell subpopulations under the different metabolic conditions. A bar graph shows the average z-score expression levels from the heatmap; Mean+/- SEM; expression levels analyzed by two-way t-tests; *p<0.01, ***p<0.001 and ****p<0.0001;

With insulin resistance, both the *Cdkn1a*^+^ and *Cdkn2a*^+^ senescent β-cell subpopulations increased (**Fig. 1C** and **Suppl. Fig. 3**). The *Cdkn1a^+^* subpopulation decreased during recovery (**Fig. 1C**), while the number of *Cdkn2a^+^* cells continued to increase (**Suppl. Fig. 3**), indicating distinct senescent populations. Principal Component Analysis (PCA) of the z-scores showed significant clustering by treatment (**Suppl. Fig. 4**); therefore, the results are shown for all three treatments in each of the senescent subpopulations.*Cdkn1a^+^* β-cells had downregulated expression of hallmark β-cell genes, including *Ins1, Ins2, Neurod1, MafA, and Pdx1*, as well as functional genes under all metabolic conditions **(Fig. 1D)**. Interestingly, this set of functional genes was not downregulated in senescent *Cdkn2a^+^* β-cells **(Fig. 1D)**, revealing functional differences between subpopulations of senescent β-cells and reconciling previous reports of increased functionality after *p16^Ink4a^* (*Cdkn2a*) overexpression in β-cells [11]. Distinct *Cdkn1a*^+^ and *Cdkn2a*^+^ subpopulations have also been identified in other tissues, including adipose tissue, liver, and heart [14].

To determine whether the upregulation of *Cdkn1a* was a transitory stress response instead of senescence, we examined co-expression with other senescence markers in various β-cell subpopulations (**Suppl. Fig. 5**). The *Cdkn1a^+^* subpopulation co-expressed *Jun* and the SASP factors *Il6* and *Il1b* while the *Cdkn2a^+^* subpopulation co-expressed *Glb1* and *Hmgb1*. These results support the notion that these two subpopulations are in a state of senescence and distinct from each other.

Expression of SASP factors in different senescent subpopulations included genes published in the SenMayo panel [15]. To minimize artifacts introduced by specific gene dropout, only genes with quantifiable reads in all three subpopulations were included in the analysis and reported as β-SenMayo (**Fig. 1E**). Compared with non-senescent and *Cdkn2a^+^*cells*, the Cdkn1a^+^* subpopulation had a significantly higher β-SenMayo score (**Fig. 1E)**, corroborating its senescent phenotype.

Validation of senescent cell subpopulations at the protein expression level included the development of a *p21^Cip1^-*tdTomato (*p21^Cip1^-*tdTom) reporter mouse model. *p21^Cip1^-* tdTom mice were generated by CRISPR/Cas9-mediated knock-in of a P2A-tdTomato cassette into the 3’-end of the endogenous *p21^Cip1^* gene in a bicistronic fashion, leading to the production of two different proteins, P21^CIP1^ and tdTomato, from the same transcript (**Fig. 2A**). Pancreatic islets isolated from these mice had detectable tdTom-expressing β-cells, as shown by immunohistochemistry (**Fig. 2B**) and fluorescent microscopy (**Fig. 2C, C’**).

**Figure 2.**
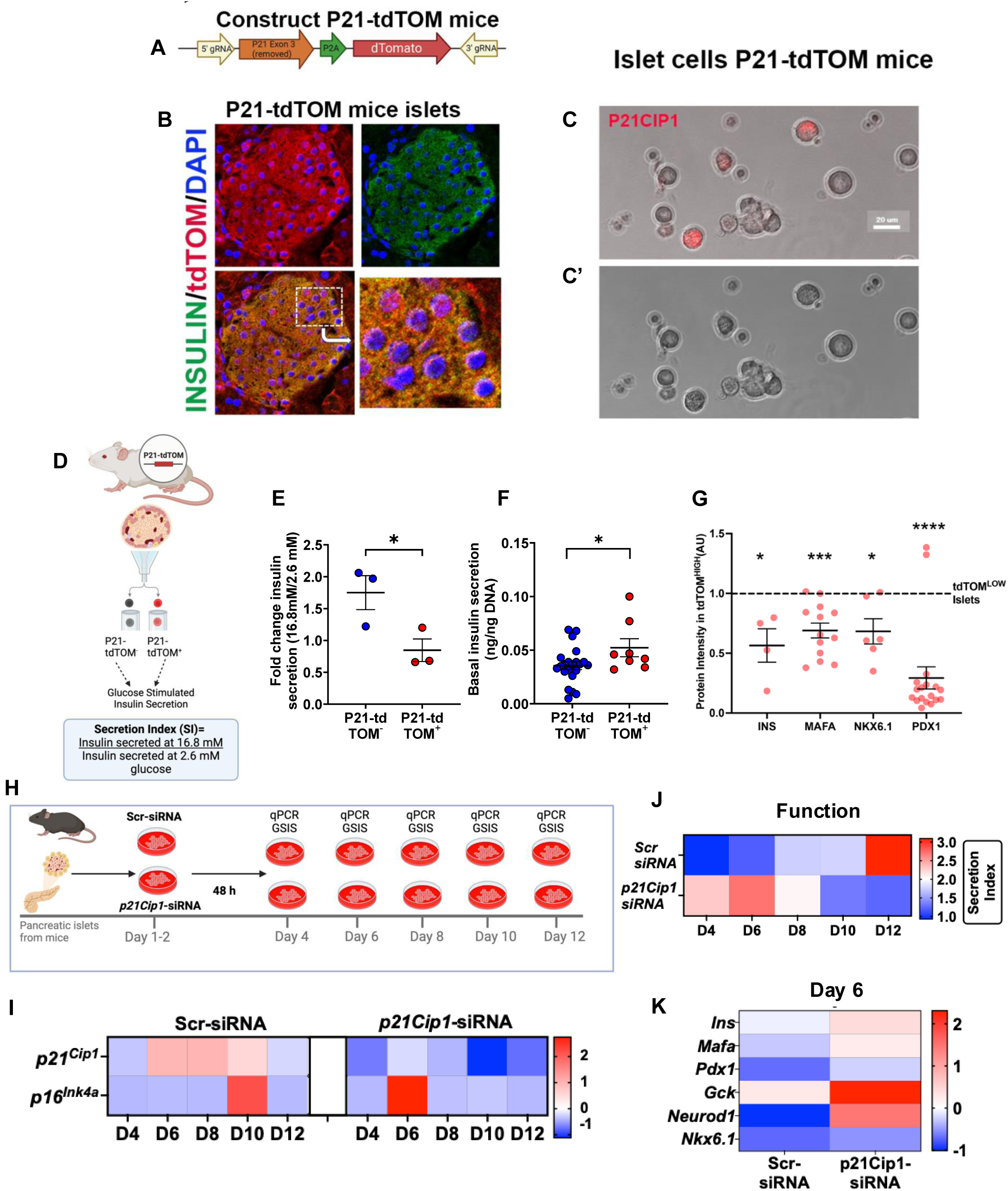
The mouse p21-Cip1-β-cell subpopulation was dysfunctional and lost its identity. **(A)** P2A-dTomato transgene used in the generation of *p21^Cip1^-*tdTomato mice, inserted at the end of the coding region using CRISPR; **(B)** Confocal microscopy reveals co-expression of insulin and *p21^Cip1^* in islet cells, confirming the presence of this senescence subpopulation. Representative image shown; **(C)** Live fluorescence with bright field microscopy of dispersed islets isolated from 3-month-old *p21^Cip^-*tdTomato, bar graph represents 20 µm **(C’)** Bright field channel only; **(D)** Islet isolation, dissociation and sorting based on tdTomato signal to purify P21-tdTOM+ and P21-tdTOM-for functional analysis; **(E)** Secretion Index= Ratio of insulin secreted at 16.8 mM glucose compared to 2.8 mM glucose, reflecting glucose responsiveness. n=3 independent animal samples; **(F)** Basal insulin secretion in P21-tdTOM+ and P21-tdTOM-cells separated by flow cytometry; n=2-5 replicates from 3 independent samples. Mean +/- SEM; islets isolated from 4 Male mice (53-70 weeks) and 3 Female mice (51-72 weeks). *p<0.05 by two-way unpaired t-test. **(G)** Loss of β-cell identity at the protein level. Quantification of protein intensity in confocal immunofluorescent images from tdTOM^HIGH^ islets respect to tdTOM^LOW^. Each point represents an individual islet from three P21-tdTOM+ mice. **(H)** Conditional p21Cip1 knock down timecourse on mouse islets to assess effects on transcription and function. (**I)** Timecourse expression levels of *p21Cip1* and *p16Ink4a* mRNA under control (Scr-siRNA) and conditional p21Cip1 knock down (p21Cip1-SiRNA, 50nM); **(J)** Timecourse of β-cell function as reflected by secretion index at different days after conditional knockdown of p21Cip1; **(K)** β-cells hallmark and function genes after *p21* conditional knockdown (p21Cip1-siRNA) at day 6. 100 islets per condition from retired male breeders n= 7 independent experiments; data shown as heatmap of z-score for panels I, K and of secretion index for J.

To test the age dependency of the P21^CIP1^ subpopulation, islets from young (3 months old) and aged (18 months old) *p21^Cip1^-*tdTomato mice were dispersed into single cells. p16^Ink4a^ was detected by immunostaining and p21^Cip1*+*^ was detected by endogenous fluorescence. All three subpopulations detected in the scRNAseq data also were detected at the protein level in islets from 3-month-old mice (**Suppl. Fig 6**) and in the following proportions: 18% P21^CIP1*+*^ and 4% P16^INK4A*+*^, supporting the scRNASeq results. The P21^CIP1+^ subpopulation significantly increased with age to 28%.

We evaluated the function of the *p21^Cip1+^* subpopulation by isolating and dispersing islets from aged adult male and female *p21^Cip1^-*tdTomato mice (51-72 weeks of age). Flow cytometry was used to separate cells into *p21^Cip1^*-tdTOM*^-^* and *p21^Cip1^*-tdTOM^+^ subpopulations, and the response to glucose was evaluated with glucose-stimulated insulin secretion (GSIS) (**Fig. 2D**). Senescent *p21^Cip1^*-tdTOM*^+^*cells had impaired glucose responsiveness, as shown by a decreased insulin secretion index (insulin secretion at 16.8 mM glucose/insulin secretion at 2.6 mM glucose) (**Fig. 2E**) due to increased basal insulin secretion (**Fig. 2F**), a feature typical of dysfunctional β-cells [2, 3, 16].

To assess whether the decreased expression of genes related to β-cell identity and function observed in the *Cdkn1a*^+^ subpopulation was reflected at the protein level, pancreatic sections from *p21^Cip1^-*tdTomato mice were stained for the following antigens and their intensity quantified: INSULIN, MAFA, PDX1, and NKX6.1. The results showed a significant decrease in the protein levels of INS, MAFA, NKX6.1, and PDX1 in islets with high levels of tdTOM compared with islets with low levels of tdTOM, as assessed by semi-quantitative immunofluorescence (**Fig. 2G and Suppl. Fig. 7**), confirming loss of cellular identity.

To evaluate the temporal patterns of *p21^Cip1+^* and *p16^Ink4a+^* expression in β-cells, mouse pancreatic islets were isolated, and a time course expression was delineated during 12 days in culture (**Fig. 2H**). In control conditions (**Fig. 2I**), *p21^Cip1^* increased from days 6-10 while *p16^Ink4a^* peaked later, at day 10. When *p21^Cip1^*-siRNA was used to keep levels down, the *p16^Ink4a^* peak occurred earlier, at day 6 (**Fig. 2I**). This timeframe is consistent with loss of identity and function in *p21^Cip1^*^+^ β-cells, since the conditional knockdown with siRNA improved β-cell function (**Fig. 2J)** and increased expression of key β -cell genes at Day 6 (**Fig. 2K**).

In summary, these analyses identified a *Cdkn1a^+^* (encoding *p21^Cip1^*) β-cell subpopulation that exhibited low glucose responsiveness, loss of β-cell identity, and increased expression of well-known senescence and SASP genes.

### Human P21^CIP1+^ senescent β-cells are dysfunctional and lose transcriptional identity revealed by spatial transcriptomics and proteomics

Analysis of the relevance of the *P21^CIP1^*^+^ β-cell population to human islets and disease included data from the Human Islets Database [17]. CDKN1A transcript levels were significantly increased in islets from donors with type 2 diabetes (T2D) compared to islets from donors without diabetes (**Fig. 3A**). Additionally, there was a negative correlation between P21^CIP1^ protein levels and insulin secretion index (**Fig. 3B**), suggesting that human P21^CIP1+^ β-cells are also dysfunctional.

**Figure 3.**
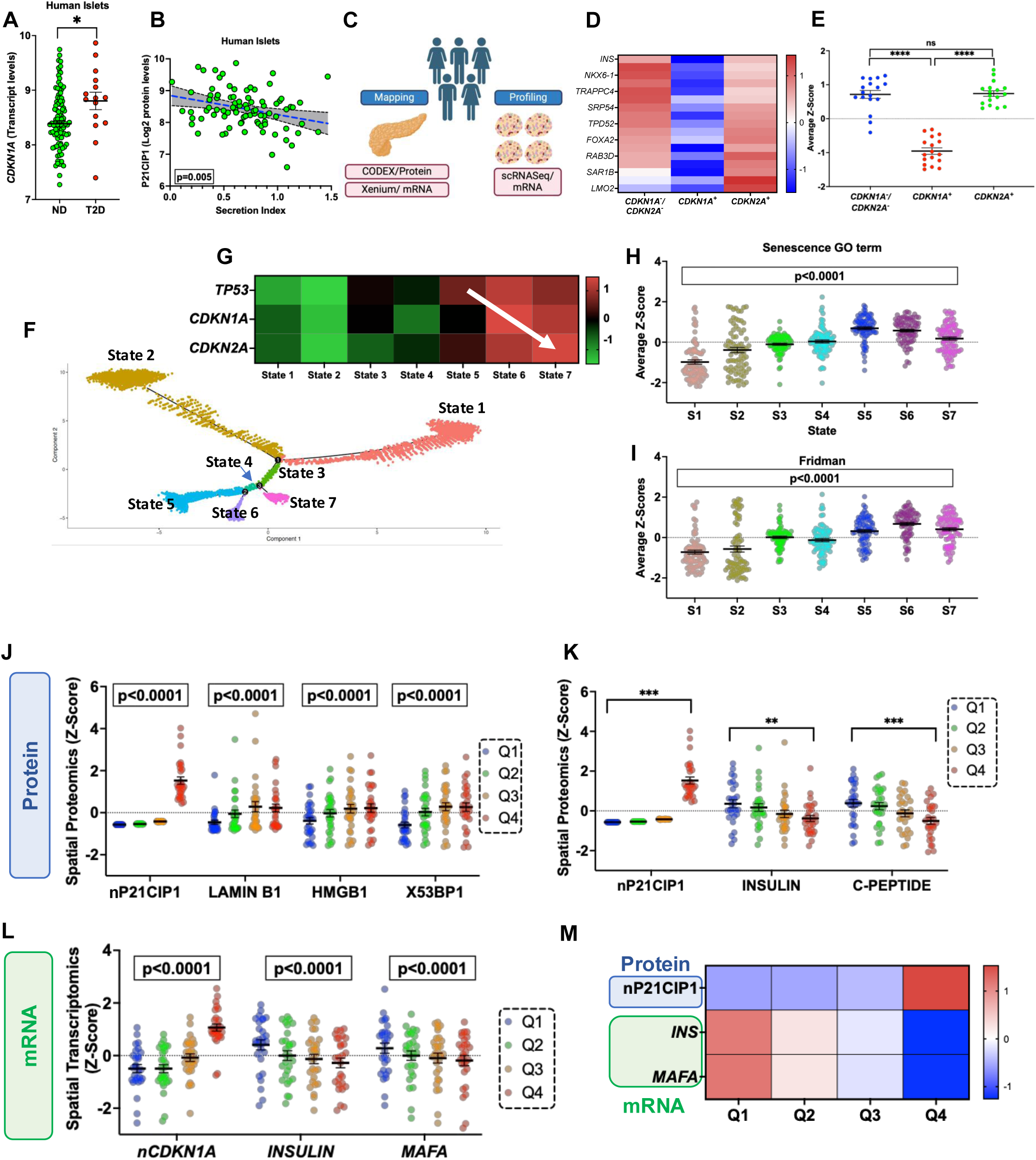
Loss of function and identity in human p21-Cip1+ β-cells. **(A)** *CDKN1A* mRNA levels in islets from human donors without (ND) and with Type 2 Diabetes (T2D). **(B)** Linear correlation between P21CIP1 protein levels in human islets and their function as expressed by the secretion index. (**C**) KAPPSen study design aimed at mapping and profiling senescent cells in both whole and dispersed human pancreas. (**D-E**) Heatmap and dot plot of human β-cell scRNASeq data for selected β-cell functional genes across three cell subpopulations: non-senescent β-cells (*CDKN1A^-^/CDKN2A^-^*), *CDKN1A^+,^* and *CDKN2A^+^;* (**F**) Pseudotime trajectory analysis of scRNASeq from non-senescent to senescent human β-cells; (**G**) Heatmap showing expression levels of key senescence genes at various stages during the senescence trajectory; (**H**) scRNASeq expression levels of genes in the senescence Gene Ontology (GO) category and (**I**) Fridman gene scores. Data shows individual cells in each trajectory state, Mean ± SEM; expression levels analyzed via 2-way ANOVA. Quartiles of nuclear P21CIP1 protein expression derived from proteomic (CODEX) data, available only from β-cells, were divided into quartiles. (**J**) Co-expression of senescence proteins within cells exhibiting quartile distribution of nuclear P21CIP1. (**K**) Co-expression of β-cell proteins within cells exhibiting quartile distribution of nuclear P21CIP1. (**L**) Quartiles of nuclear CDKN1A mRNA expression from Xenium and co-expression with hallmark β-cell genes. (**M**) Heatmap showing overlapping co-expression of nuclear P21CIP1 with β-cell hallmark genes.

To further profile and map *P21^CIP1^*^+^ β-cells in human islets, whole pancreas and human islets were obtained as part of the KAPP-Sen Tissue Mapping Center (**Fig. 3C**). Isolated islets from donors without diabetes across the lifespan underwent scRNASeq for transcriptomic analysis. Key β-cell identity and functional genes (*INS, NKX6.1*) were significantly downregulated in the *CDKN1A^+^* β-cell subpopulation when compared to non-senescent and *CDKN2A^+^* β-cells (**Fig. 3 D, E**). To understand whether these senescent subpopulations may have originated from independent lineages as opposed to a single population that expresses these markers in a temporally distinct fashion, as previously suggested by us and others [6, 18, 19], pseudotime analysis of scRNASeq was performed in human β-cells. Seven states were identified during the progression of a human non-senescent β-cell to a senescent state (**Fig. 3F, G**), with states 5, 6, and 7 showing marked upregulation of key senescent genes: *TP53, CDKN1A*, and *CDKN2A* (**Fig. 3G**). Interestingly, state 5 expressed higher levels of *TP53*, state 6 of *CDKN1A* and state 7 of *CDKN2A*, suggesting a temporal progression of β-cells through these three senescent stages. In support of these being senescent states, 5, 6, and 7 had significantly higher senescence scores using the GO term (**Fig. 3H**) and Fridman (**Fig. 3I**) gene data sets.

Given the correlation between P21^CIP1^ with T2D and islet dysfunction in humans, spatial proteomic (CODEX) and transcriptomic (Xenium) analysis focused on CDKN1A/P21^CIP1+^ β-cells. P21^CIP1^ expression was divided into quartiles (Q), with Q1 having the lowest levels and Q4 the highest. At the protein level, β-cells with higher levels of P21^CIP1^ also had higher levels of other senescence markers: LAMIN B1, HMGB1, and XP53BP1 (**Fig. 3J**). Consistent with their loss of identity, this same population had lower levels of key β-cell proteins: INSULIN and C-PEPTIDE (**Fig. 3K**). At the transcriptomic level, β-cells with the highest levels of nuclear *CDKN1A* had the lowest levels of *INSULIN* and *MAFA*, a key functional transcription factor (**Fig. 3L**). These correlations were maintained when β-cells were selected based on nuclear protein P21^CIP1^ levels and their transcriptional mapping revealed lower levels of *INSULIN* and *MAFA* mRNA transcripts (**Fig. 3M**).

These results demonstrate that a senescent and dysfunctional subpopulation of P21^CIP1+^ β-cells can be identified in human islets. These cells appear to be more abundant in tissues from individuals with T2D, and thus may contribute to β-cell dysfunction and T2D pathogenesis.

### Non-cell autonomous effects of the **β**-cell SASP include secondary senescence

To assess whether SASP factors from senescent β-cells had non-autonomous effects on non-senescent β-cells and contribute to senescence, a series of experiments were conducted using complete conditioned media (CM) containing the full SASP from senescent cells, a selection of 4 SASP factors and individual SASP factors.

For complete CM experiments, a mouse insulinoma cell line, (MIN6) was treated with CM from senescent and non-senescent cells. CM from senescent cells was generated by treating MIN6 cells, with the DNA-damaging agent, bleomycin, which induces senescence and SASP secretion *in vitro* [6]. Conditioned media from vehicle-treated cells (CCM) or bleomycin treated cells (BCM), were collected and used to treat naïve MIN6 cells for 24 h. Five days after exposure to CCM or BCM, cells were collected and expression of senescence markers and proliferation were analyzed **(Fig. 4A)**. Exposure to BCM resulted in increased *p16^Ink4a^*expression **(Fig. 4B)** and a marginally increased β-gal activity **(Fig. 4C)**, both of which are senescence markers. These findings support the non-cell autonomous effects of the SASP, capable of inducing secondary senescence.

**Figure 4.**
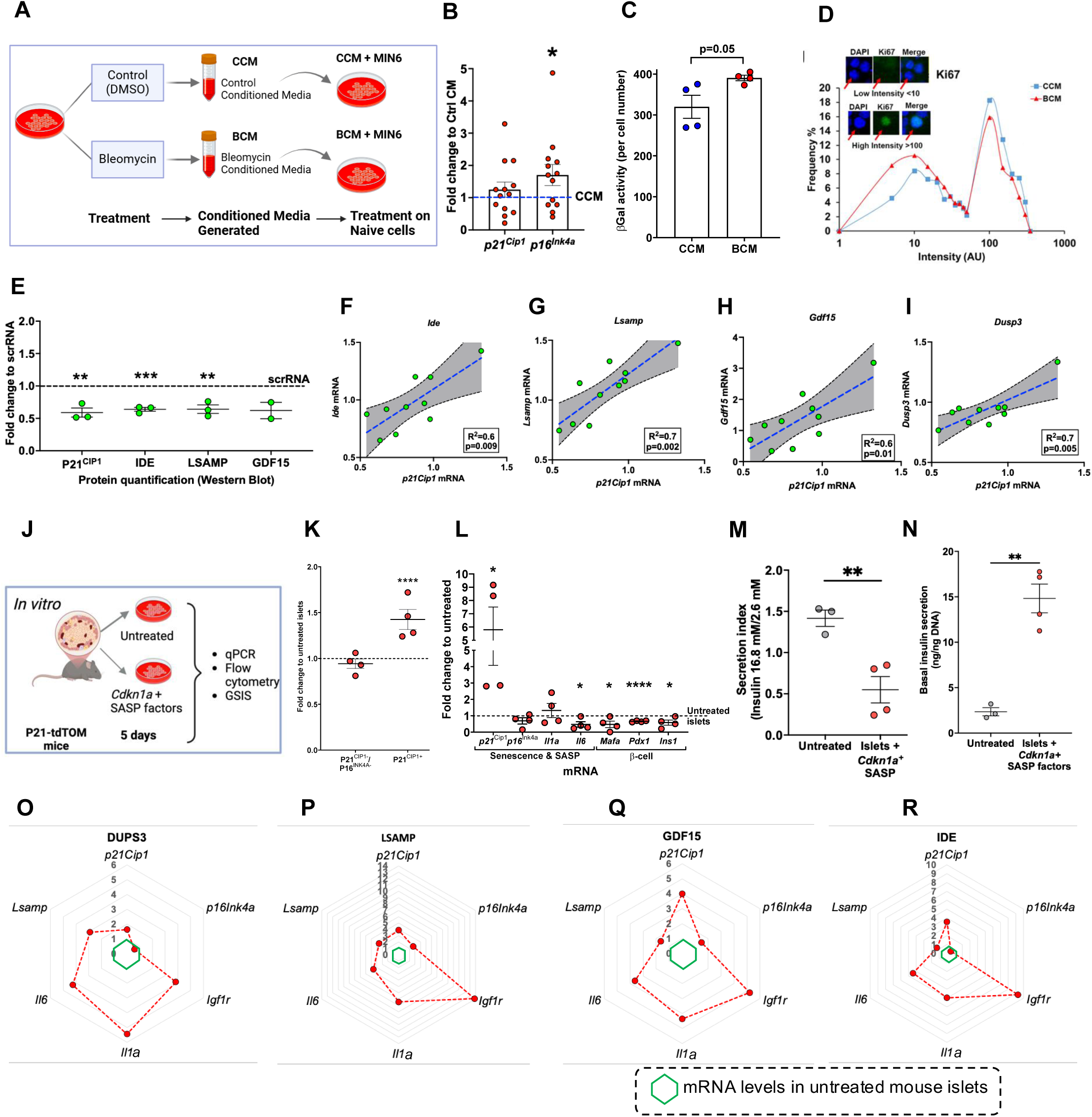
Secondary senescence: β-cell SASP has non-cell autonomous effects on neighboring cells. **(A)** Workflow to test the effects of SASP on naïve MIN6 cells. **(B)** Effects of BCM on *p21^Cip1^* and *p16^Ink4a^* mRNA expression of naïve MIN6 cells (n=13-14 individual samples from 3 independent experiments*) *p<0.0 5*, analyzed by non-parametric Wilcoxon test; **(C)** β-Gal^+^ activity normalized per cell number (n=4 individual samples); **(D)** Cumulative frequency graph of Ki67 staining of MIN6 cells reflecting proliferating subpopulations (Cells counted: n=2911 for CCM (4 individual samples from two different experiments), n=3000 for BCM (4 individual samples from two different experiments)); **(E)** Effects of conditional knockdown of *p21Cip1* achieved using siRNA (50 nM). Protein quantification by Western Blot of selected Cdkn1a+ SASP factors after treatment with P21Cip1 siRNA at 50 nM in MIN6 cells (n=3 independent experiments). Mean +/- SEM **p<0.005 and ***p<0.0005; **(F-I**) Expression levels of *Cdkn1a*-SASP factors from individual samples and their correlations with *Cdkn1a* expression. Lines of best fit are shown, along with dotted lines indicating 95% confidence intervals. P-values were calculated using the null hypothesis that the slope of the best fit line equals 0; **(J)** Treatment of islets from P21-tdTom mice with Cdkn1a SASP factors (LSAMP+DUSP3+GDF15+IDE) during 5 days; **(K)** Cellular subpopulations as determined by flow cytometry of dispersed islets after treatment; **(L)** Effects of *Cdkn1a+* SASP factors on transcription of selected senescence and key β-cell genes; **(M)** Secretion index of islets treated with *Cdkn1a+* SASP factors; **(N)** Basal insulin secretion (n-4 independent experiments). Mean+/- SEM; expression levels analyzed by two-way t-tests; *p<0.01, **p<0.001 ***p<0.0001 and ****p<0.00001. **(O-R)** Radar plots of qPCR results for the mean expression of senescence and SASP genes. MIN6 cells treated with an individual selected SASP protein for 48 hours, followed by a further 48-hour incubation. Mean+/-SEM; n= 2 independent experiments with 2-3 replicates per experiment. Concentrations of SASP proteins used in culture media are shown in **Table 1** and are based on human circulating concentrations of SASP factors. Mean+/- SEM; expression levels analyzed by two-way t-tests; *p<0.01, **p<0.001 ***p<0.0001 and ****p<0.00001. The midpoint of 0 represents average expression across all samples.

**Table 1.**
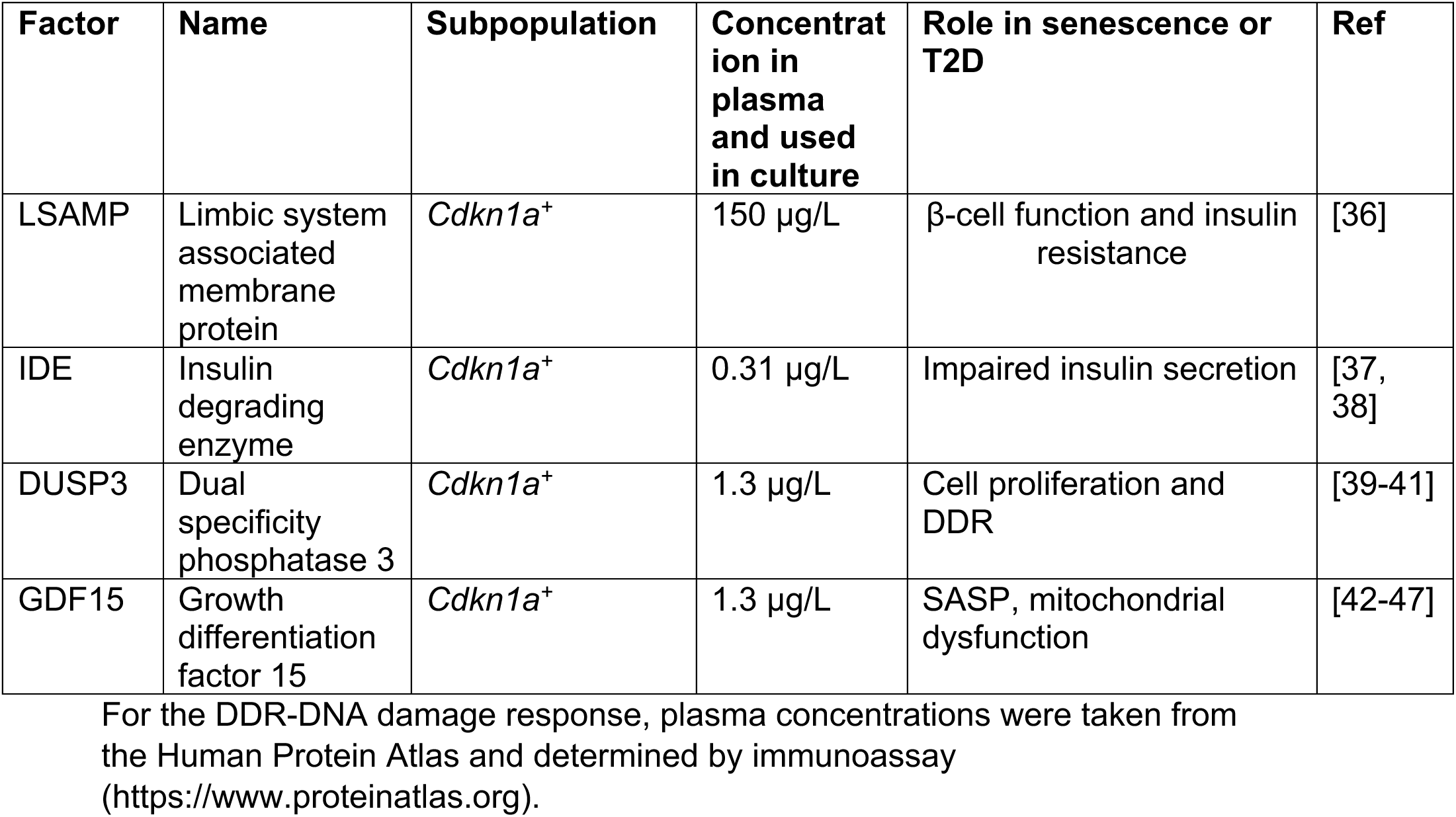
Selected SASP factors and their role according to senescence cell subpopulation.

Proliferative arrest, an additional indicator of senescence, was analyzed by measuring Ki67 staining intensity in individual nuclei using ImageJ. A frequency distribution graph of Ki67 intensity revealed two main subpopulations: non-proliferative cells (*Ki67* intensity <10 AU, as measured with image analysis software) and proliferative cells (Ki67 intensity>100 AU) **(Fig. 4D)**. MIN6 cells treated with conditioned media obtained from cells previously treated with bleomycin (BCM), had a greater percentage of nonproliferating cells and a lower percentage of proliferating cells, suggesting induction of senescence.

To analyze the non-cell autonomous effects of individual SASP factors in β-cells, four factors were selected based on the following criteria: 1) previously published as a senolytic target and/or SASP factor; 2) identified as a SASP factor secreted by β-cells in our proteomic analysis [6]; 3) common factor between mouse and human SASP and/or reported increase in β-cells from donors with T2D [6]; and 4) measured concentration in plasma by immunoassay as reported in the Human Protein Atlas (**Table 1 and Suppl. Table 2**). The following SASP factors met these criteria and were preferentially transcribed by *Cdkn1a^+^* cells: LSAMP, IDE, DUSP3, GDF15.

To test the dependency of these SASP factors on *p21^Cip1^*, a conditional siRNA knockdown was developed for *p21^Cip1^*. At the protein level, conditional downregulation of *p21^Cip1^* led to significant decreases in IDE and LSAMP (**Fig. 4E**). At the transcriptional level, a significant positive correlation was observed between *Lsamp*, *Dusp3*, *Gdf15*, and *Ide,* and the mRNA levels of *p21^Cip1^* (**Fig. 4F-I**). These patterns show that the chosen SASP factors directly correlated with the CDK inhibitor, *p21^Cip1^*, thereby supporting subpopulation specificity.

To test the non-cell autonomous effects of *p21^Cip1^*-SASP factors, islets were isolated from male and female *p21*-tdTOM mice and incubated with a combination of *Cdkn1a*^+^ SASP factors (LSAMP, DUSP3, GDF15, and IDE) for 5 days before assessing senescence and function (**Fig. 4J**). Protein concentration levels were determined by reported circulating levels in humans in the Human Protein Atlas, however, the paracrine concentrations are likely higher than those reported in plasma but impossible to measure with currently available technology. SASP from *Cdkn1a*^+^ cells significantly increased the P21^CIP1+^ subpopulation as measured by flow cytometry (**Fig. 4K**) and upregulated *p21^Cip1^* mRNA (**Fig. 4L**) while downregulating the key β-cell genes: *Mafa, Pdx1*, and *Ins1*. At the functional level, *Cdkn1a^+^* SASP factors impaired β-cell function, as reflected by a lack of glucose responsiveness (**Fig. 4M**), characterized by increased basal insulin secretion (**Fig. 4N**).

To test whether individual *Cdkn1a^+^* SASP factors were able to induce secondary senescence, MIN6 cells were treated with an individual protein for 4 days. All four proteins upregulated expression of 6 senescence and SASP genes (*p21Cip1, p16Ink4a, Igf1r, Il1a, Il6, and Lsamp*) (**Fig. 4O-R)**; LSAMP was the individual factor that induced the greatest increases in gene expression (**Fig. 4P**).

These results indicate that SASP factors induced secondary senescence in islet cells in a pathway-specific way: SASP released from p21^Cip1+^ cells increased the p21^Cip1^ subpopulation and gene transcription. SASP factors induced senescence both in combination and individually, implying that a small percentage of senescent p21^Cip1+^ β-cells can induce secondary senescence and loss of β-cell identity. Based on these findings, the β-cell SASP can be considered an additional therapeutic target for restoring β-cell identity and functionality.

### JAK inhibitors restored β-cell function, transcriptional identity, and cell subpopulations in mouse islets

SASP secretion and action are known to be regulated by several pathways [20] whose regulation was queried in mouse senescent β-cells [3]. A significant and prominent upregulation of the JAK/STAT pathway (**Fig. 5A**) was identified compared with non-senescent β-cells, and JAK inhibitors were tested *in vitro* and *in vivo* (**Fig. 5B**) to assess their efficacy in maintaining β-cell function and identity while decreasing senescence.

**Figure 5.**
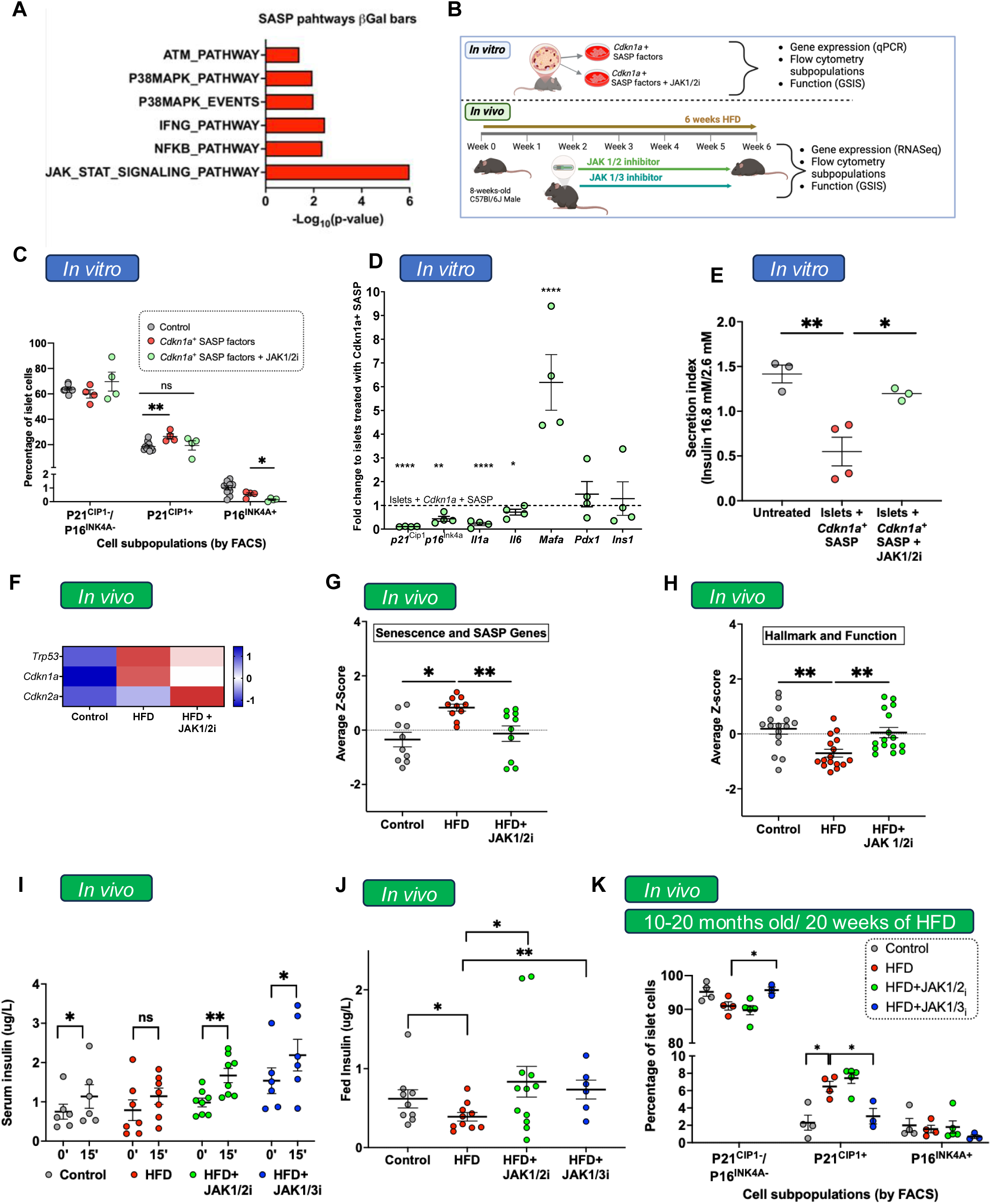
Secondary senescence induced by *Cdkn1a+* SASP factors is counteracted by JAK/STAT inhibitors in mouse islets in vitro and in vivo. **(A)** Pathway analysis of SASP-regulating pathways in RNA-seq data from mouse senescent β-cells (3) reveals upregulation of the JAK/STAT pathway. **(B**) Experimental design *In vitro:* Treatment of islets from P21-tdTom mice with Cdkn1a SASP factors (LSAMP+DUSP3+GDF15+IDE) in combination with JAK1/2 inhibitor (momelotinib). *In vivo:* male C57Bl/6 mice were fed a high-fat diet (HFD) for 6 weeks and treated for 4 weeks with either tofacitinib (JAK1/3 inhibitor) or momelotinib. n=9 control, n= 11 HFD group,; n= 13 HFD + momelotinib and n=6 HFD + tofacitinib; **(C)** Flow cytometry analysis of dispersed islets treated with Cdkn1a SASP factors with or without JAK1/2 inhibitors (JAK1/2i); **(D)** Transcriptional analysis of islets treated with Cdkn1a SASP factors with or without JAK1/2i; **(E)** Functional analysis of islets treated with Cdkn1a SASP factors with or without JAK1/2i. n=4 independent experiments. Mean+/- SEM; expression levels analyzed by two-way t-tests respect to control; *p<0.01, **p<0.001 ***p<0.0001 and ****p<0.00001. **(F)** Heatmap showing expression of key senescent genes in islets isolated from different treated groups; **(G)** Average z-score expression of senescence and SASP genes; **(H)** Average z-score expression of β-cell hallmark and functional genes, mean z-scores per group are shown along with their average+/- SEM. Expression levels were analyzed by two-way ANOVA followed by a Fridman test; **p<0.001, ****p<0.00001. **(I)** Glucose stimulated insulin secretion by sampling plasma at minutes 0 and 15 during IPGTT. (**J**) Fed plasma insulin in mice. Mean +/- SEM; *p<0.01, **p<0.001 by two-way unpaired t-test. **(K)** Subpopulations of senescent islet cells analyzed by flow cytometry in islets from treated and untreated mice. Female C57BL/6J mice, aged 10-20 months, were fed a HFD for 20 weeks and treated with momelotinib or tofacitinib at doses used in human clinical trials (Momelotinib: 1.5 mg/kg, Tofacitinib: 1 mg/kg). n=5 HFD+ Momelotinib (42-83 weeks of age), n=4 HFD+ Tofacitinib (42-68 weeks of age), n=4 HFD (48-83 weeks of age), n= 4 control (42-83 weeks of age). Mean +/- SEM; *p<0.05 by two-way unpaired t-test.

*In vitro*, JAK1/2i (momelotinib) attenuated the effects of *Cdkn1a*-SASP factors by decreasing the p21^CIP1^ and P16^INK4A^ subpopulations of senescent cells measured by flow cytometry in islets treated with *Cdkn1a* SASP factors (**Fig. 5C**). At the transcriptional level, JAK1/2i decreased all tested senescence and SASP gene expression in islets treated with *Cdkn1a^+^* SASP factors (**Fig. 5D**), while upregulating *Mafa*, a key transcription factor for β-cell function. These positive transcriptional and subpopulation changes were reflected at the functional level. Concurrent treatment of mouse islets with SASP factors and JAK1/2i restored the secretion index, indicating improved β-cell functionality (**Fig. 5E**).

The effects of JAK inhibitors were also evaluated *in vivo* with a model of insulin resistance in which mice were fed a high-fat diet (HFD) that increases senescence in β-cells [3] and other metabolically relevant tissues [21]. Young (8-week-old) *C57Bl/6J* male mice were fed a HFD. They received an osmotic pump containing vehicle, JAK1/2 inhibitor (momelotinib), or a JAK1/3 inhibitor (tofacitinib) for the last 4 weeks (**Fig. 5B**). Bulk RNA-Seq of islets isolated from animals in the different treatment groups confirmed *Cdkn1a* induction by HFD (**Fig. 5F**), increased expression of senescence and SASP genes (**Fig. 5G)**, and decreased expression of β-cell hallmark identity and functional genes (**Fig. 5H**). Downregulation of the JAK pathway was confirmed at a transcriptional level (p<0.02), with no differences in the SMAD/Activin pathway, which can be modulated by JAKi. At the functional level, both JAK1/2i and JAK1/3i restored glucose responsiveness lost after HFD *in vivo* evaluated during an intraperitoneal glucose tolerance test (IPGTT) (**Fig. 5I**) and increased circulating insulin levels (**Fig. 5J**).

The chronic effects of the JAK inhibitors were evaluated over 20 weeks in middle-aged (6-9 months old) male mice, at a dose similar to that used in humans (momelotinib 1.5 mg/kg, tofacitinib: 1 mg/kg, oral administration, 5 days per week throughout the whole study). The selection of age and duration was based on the known increase of senescent β-cells in islets from mice with chronic insulin resistance and older age. At the end of the study, islets were isolated, and the subpopulations of senescent cells were analyzed by flow cytometry (gating strategy is shown in **Suppl. Fig. 8**). JAK1/3i increased the percentage of the non-senescent subpopulation while decreasing the percentage of the P21^CIP+^ subpopulation that was induced by HFD (**Fig. 5K**), consistent with therapeutic effectiveness.

The peripheral effects of JAK1/2i were evaluated using qPCR in the following metabolically relevant tissues: liver, visceral adipose tissue, and skeletal muscle (**Suppl. Fig. 9**). JAK 1/2i did not seem to have a significant effect on expression of senescence or SASP genes at the doses used (12.5 mg/kg/day), suggesting a preferential effect on β-cells in the model used.

These results showed that JAK inhibitors decrease senescence genes and subpopulations while restoring β-cell function and transcriptional identity in mouse β-cells, both *in vitro* and *in vivo*.

### Pharmacological JAK inhibition restored function of human β-cells

To analyze the relevance of JAK inhibition in human β-cells as a potential regulator of secondary senescence, data from the Human Islet Program was used [17]. JAK1 protein levels were increased in islets from donors with T2D compared to islets from donors without diabetes (**Fig. 6A**). Additionally, there was an inversely negative correlation between JAK1 protein levels and secretion index (**Fig. 6B**), suggesting that human JAK1 might be a potential target to restore β-cell function in islets with higher levels of P21^CIP1+^. Based on this rationale, JAK1/2i was tested in human islets.

**Figure 6.**
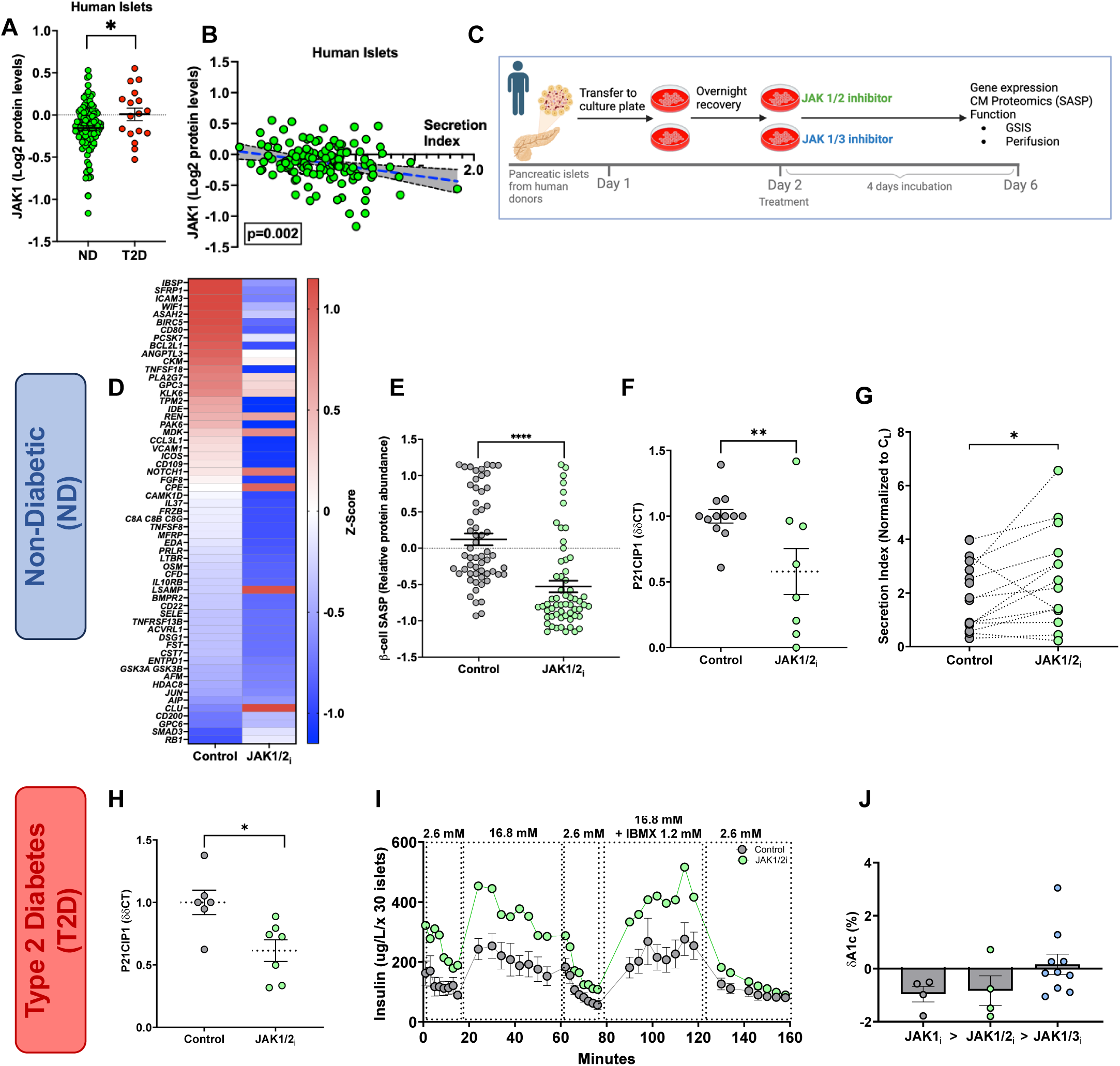
Effects of JAK1/2i on secretion of human β-cell function. **(A)** JAK1 protein levels in human islets from non-diabetics (ND) and Type 2 diabetes (T2D) donors**; (B)** linear regression of JAK1 protein levels and secretion index in from human islets; (**C**) Workflow of human islets from ND and T2D donors treated with JAK1/2i or JAK1/3i for 4 days. (**D-E**) Heatmap and dotplot of SASP (log_2_-protein abundance) secretion in conditioned media collected from human untreated (control) or treated (JAK1/2i). Top upregulated human β-cell SASP proteins were selected. Conditioned media was collected as specified from 5 young non diabetic human donors (**Suppl. Table 3**) and a paired analysis per analyte was performed between conditions. Cell number varied from one donor to another but was maintained constant across different treatments. Donor 1-526,000 cells/treatment; donor 2-192,000 cells/treatment; donor 3-231,000 cells/treatment; donors 4&5-87,500 cells/treatment. (**F**) P21CIP1 mRNA levels in human islets from ND donors. (G) β-cell function evaluated by GSIS in human islets from ND donors (Donor information in **Suppl. Table 3**) with 1-3 replicates per donor. (**H**) Effects of JAK1/2i in P21CIP1 mRNA in islets from T2D donors; (**I**) β-cell function evaluated with islet perifusion in islets from a donor with T2D, BMI 38, 58 years old. **(Suppl. Table 3)**. **(J)** Mean A1c change before and after starting treatment with a specified JAK inhibitor in people with Type 2 Diabetes.

Islets from donors with and without T2D (**Suppl. Table 3)** were treated *in vitro* with JAK1/2i for 4 days, after which CM was collected for aptamer-based proteomic analysis of the SASP (**Fig. 6C**). The top upregulated **(Fig. 6D)** human SASP proteins identified in human β-cells [6] were compared between CM from control human cells to CM human cells treated with JAK1/2i. Treatment with JAK1/2i resulted in a significant decrease in the secretion of SASP proteins **(Fig. 6D, E),** demonstrating its effectiveness in targeting human SASP.

To analyze the effects of SASP inhibition on senescent subpopulations, islets from the donors without diabetes were treated as described above and analyzed for the expression of *P21^CIP1^* mRNA, which was decreased by JAK1/2i **(Fig. 6F)**. This was accompanied by an increase in the secretion index in islets from donors without T2D (**Fig. 6G**).

To test whether the JAK pathway was also relevant in a diseased state, islets from donors with T2D were treated with JAK1/2i for four days. *P21^CIP1^*transcription was significantly decreased (**Fig. 6H**). From a functional perspective, JAK1/2i restored insulin secretion dynamics in the islets of a donor with T2D (58 years old and BMI 38) (**Fig. 5I**). This can be seen as a restoration of first-phase insulin secretion and an enhanced response to the secretagogue, IBMX.

Several JAK inhibitors are approved for clinical use for different diseases (*e.g*., rheumatoid arthritis). Deidentified A1c levels were obtained from the Joslin Adult Clinic from a limited number of adults with T2D who were also prescribed a JAKi. Analysis of the longitudinal A1c of each patient, revealed that individuals taking JAK1i and JAK1/2i had A1c decreased by 1% compared to levels before the drug was prescribed (**Fig. 6J**).

Based on these results, we conclude that the human β-cell SASP is a therapeutic target to restore function during progression and in established T2D.

## DISCUSSION

The importance of heterogeneity in cellular senescence is increasingly being recognized as a key factor in developing better-targeted interventions for various diseases. *p21^CIP1^* (encoded by *Cdkn1a*) and *p16^INK4a^*(encoded by *Cdkn2a*) are well-established markers and effectors of senescence, and their expression patterns have been used to define three different subpopulations of β-cells. We found that p21^CIP1+^ cells are the predominant senescent subpopulation of β-cells in models of metabolic stress, T2D, and aging. Interestingly, the identity of these cells at the gene expression level varied from that of p16^INK4A+^ cells: *Cdkn1a^+^* β-cells were characterized by reduced expression of hallmark genes related to β-cell identity and function, along with an increase in commonly described SASP genes [3].

The co-expression of other markers of senescence and increased transcription of SASP factors support this is a *bona fide* senescent population rather than a population undergoing transient stress. The expression dynamics observed in islets from mice treated with S961 for two weeks suggested that *Cdkn1a* could be an initial driver of β-cell senescence, while *Cdkn2a* would increase at a later stage, a process that has also been proposed by others [19]. This concept was further supported by pseudotime analysis of human scRNASeq of β-cells, and by the conditional knockdown *of p21^Cip1^* in mouse islets. Even though these populations are temporally asynchronous, both populations arise from a common state and, therefore, can be independently regulated; their specific control should be studied further.

Our data support the induction of secondary senescence in β-cells by *Cdkn1a*^+^ cells. This effect was shown with full SASP in conditioned media, with a subset of selected SASP factors and by each factor individually. In these different conditions, naïve cells exposed to SASP exhibited increased expression of senescence genes (particularly *p21^Cip1^*) and became dysfunctional, consistent with secondary senescence being an important effector of β-cell dysfunction in models of insulin resistance and progression to T2D. The role of the β-cell SASP as an inducer of secondary senescence underscores the importance of therapeutically targeting the secretome of senescent cells through pharmacological interventions. Treatment of β-cells with the JAK1/2i decreased the expression of senescence markers and improved cell function in models of early T2D. JAK1/2i, specifically momelotinib, competes with Janus tyrosine kinases 1 and 2 to reduce the intracellular signalling of cytokines and growth factors, thereby mitigating their transcriptional effects [22, 23]. It has also been shown to inhibit the SASP profile of senescent cells [10]. In human β-cells, JAKi significantly decreased secretion of SASP factors and, based on RNASeq data, it also significantly decreased signalling by interleukins (IL2, 6, 9, 20, 21 27, 28, 35), IFNγ, and TP53 activity. The specific roles of each of these pathways in restoring the function and identity of β-cells will be subject of future studies.

In addition, preclinical trials have shown that JAK1/2i inhibits tumor growth and promotes apoptosis in ovarian, colorectal, and brain cancer [24–27]. Therefore, the reduction in *p21^CIP1^* and *p16^INK4a^* could be due to a decrease in the number of cells entering senescence rather than a direct effect on cell cycle regulation or escape from the senescent state, which decreases the potential tumorigenicity of these agents.

If JAK inhibitors are considered as an additional strategy for restoring β-cell function during progression to T2D, the reported side effects of these agents should be considered. Their main reported side effects are infections, particularly viral infections, which can be mitigated with vaccination [28]. Also increased risk of cardiovascular and thromboembolic events which could be avoided by specifically targeting these drugs to β-cells through chemical manipulation. Supporting the use of JAK inhibitors to protect human β-cells, daily administration of baricitinib to people, including children, with recent onset Type 1 Diabetes, preserved β-cell function [29], and was well tolerated.

In conclusion, senescent β-cell heterogeneity provides a biological basis for differentially targeting specific subpopulations and counteracting their effects on neighbouring cells as a contributing factor to diabetes progression. Differential pharmacological targeting strategies are available, and inhibiting the β-cell SASP with JAK inhibitors has been shown to restore human β-cell function and enhance cell survival in models of early β-cell dysfunction. These concepts further our understanding of senescence biology and support the rational development of novel therapeutic strategies for diabetes.

## MATERIALS AND METHODS

### Animals

Male *C57BL/6J* mice (8-week-old or 6-9-month-old-retired breeders) obtained from Jackson Labs were used for islet isolation and conditioned media generation. Male mice were chosen at this stage because they are more susceptible to develop insulin resistance after dietary or pharmacological interventions. All experiments were performed in the animal facilities at the Joslin Diabetes Center with approval from its Animal Care and Use Committee. Mice were kept on a 12-hr light/dark cycle with water and food provided *ad libitum*. A housing temperature of 22.2-22.7°C was maintained.

### *p21^Cip1^*-tdTomato Mice

To generate a mouse in which dTomato expression was driven by the *p21^Cip1^* gene, the P2A-dTomato transgene was inserted in place of the stop codon at the 3’ end of the coding region. The Easi-CRISPR design methodology was followed as previously published [30, 31]. Bioinformatic analysis of the gene for gRNA identification was performed by crispor.tefor.net. Microinjection was performed into embryos from 4-week-old *C57 Bl6* female donors (Jackson), which were subsequently implanted into 6-week-old *Swiss Webster* females (Charles River). Junction sequencing was performed to verify the construct.

### Cell Lines

Murine MIN6 cells were originally provided by Dr. Jun-ichi Miyazaki from Osaka University Medical School, and the sex of the cell line was not available. The cell line was maintained in Dulbecco’s modified Eagle’s medium (HG, w/4500 mg/L glucose, L-glutamine, sodium pyruvate, and sodium bicarbonate) from Sigma‒Aldrich® supplemented with 15% fetal bovine serum (FBS) from Summerlin Scientific©, 1% penicillin‒streptomycin (5,000 U/mL) from Gibco™, and 0.05% β-mercaptoethanol (99% cell culture tested) from Sigma‒Aldrich®. The cells were incubated at 37°C with 5% CO_2_.

### Mouse Primary Islets

*INK-ATTAC* and *C57BL/6J* mice were used for islet isolation. Isolation was performed according to the protocol described by Gotoh, M.*, et al.* [32]. The islets were then hand-picked under a StereoZoom® 7 microscope and plated in Petri dishes with islet media. The media were composed of RPMI 1640 (1x, [+] L-glutamine) (Gibco™) supplemented with 10% fetal bovine serum (FBS; Summerlin Scientific) and 1% penicillin‒ streptomycin (5,000 U/mL; Gibco). For the dispersion of islets, we used TrypLE™ Express Enzyme (1x, [-] Phenol red) (Gibco). The islets were incubated at 37°C with 5% CO_2_.

### Human Primary Islets

Human islets were obtained from donors through IIDP and Prodo Labs (**Suppl. Table 3**). Immediately upon arrival, the islets were cultured in CMRL Medium 1066 (1x, [-] glutamine) from Gibco™ containing 10% FBS (Summerlin Scientific) and 1% penicillin‒ streptomycin (5,000 U/mL) (Gibco). For the dispersion of islets, we used TrypLE™ Express Enzyme (1x, [-] Phenol red) (Gibco). The islets were incubated at 37°C with 5% CO_2_.

### Whole Pancreas Preparation

All the whole pancreases were procured by the University of Texas Health San Antonio from IRB-exempt brain-dead donors and then shipped to Joslin Diabetes Center, Harvard Medical School. Inclusion criteria: Age range 20-60+. Exclusion criteria: type 2 diabetes, pancreatic malignancy, pancreatitis, use of agents with senolytic activity. The pancreases were cut into the head, body, and tail of the locations along the pancreas carcinoma guideline, cut into 4mm thickness, and placed into order paraffin or frozen cassettes. The paraffin cassettes were fixed in 4% paraformaldehyde for 24 hrs and processed for paraffin embedding at Joslin Histology Core. The frozen cassettes were molded with O.C.T. compound (Fisher Scientific No.23-730-571) and frozen in the cold 2-methylbutane in a beaker surrounded by dry ice and 100% ethanol (Fisher Scientific No.60-048-072) bath followed by drying and storage at -80°C. At least three paraffin cassettes per each location of head, body, and tail were shipped to The Jackson Laboratory; all the procedures were done at 4°C and avoided exposure to direct light. A detailed protocol can be found in https://www.protocols.io/view/kapp-sen-tmc-whole-pancreas-preparation-4r3l224nql1y/v1?step=1

### JAK Inhibitor Treatment *in vitro*

The following drug concentrations were used for treatment and were diluted in CMRL Medium 1066 1X (See Human Primary Islets): 1 µM JAK1/2i (momelotinib), and 1 µM JAK1/3i (tofacitinib). The concentrations of momelotinib and tofacitinib were based on published data [10], which showed effective decreases in the transcription of specific SASP factors. Human and mouse cells were incubated for 4 days in wells previously treated with PEI and Geltrex.

### JAK Inhibitor Treatment *in vivo*

After two weeks of HFD feeding (60% v/v), the animals (8-week-old or 6-9-month-old male *C57BL/6* mice) were subdivided into four groups: saline, JAK1/3i tofacitinib, JAK1/2i momelotinib, vehicle (HFD, 0.9% NaCl), JAK1/3i (HFD+tofacitinib, 10 mg/kg/day), or JAK1/2i (HFD+momelotinib, 12.5 mg/kg/day) administered through through an osmotic pump in 8-week old animals. All control animals received vehicle via oral administration (NaCl 0.9%). Blood glucose levels in the mice in all groups were monitored biweekly, and the mice were sacrificed after treatment (4 weeks for HFD). The pancreatic islets were then isolated for qPCR analysis or bulk RNA-seq. Heart blood was collected through cardiac puncture to measure circulating insulin in the mice. For the 20-week HFD trial, doses of JAK STAT inhibitors were modified as follows to match those used in human clinical trials: JAK1/2i, 1.5 mg/kg; and JAK1/3i, 1 mg/kg, they were administered through oral gavage 5 days/week throughout the length of the study.

### Intraperitoneal Glucose Tolerance Test (IPGTT)

The IPGTT was performed before and after senolytic and senomorphic treatment. Mice fasted for 6 hours received an intraperitoneal injection of glucose (2 g/kg body weight). Blood glucose was measured at 0 (baseline before injection), 15, 30, 60, 90, and 120 minutes post-injection. Blood from the tail was collected at 0 and 15 minutes for serum analysis of insulin via ELISA (Mercodia 10-1247-10).

### Generation of SASP-Conditioned Media

MIN6 cells were passaged and plated into two separate Nunc™ EasYFlask™ Cell Culture Flasks from Thermo Scientific™ with 10 mL of normal media (see Cell Lines) using approximately 7,000,000-10,000,000 cells per flask. The cells were cultured for 24 hrs to allow attachment and incubated at 37°C with 5% CO_2_. The flasks then had their normal media replaced with media containing either 50 μM bleomycin sulfate (ApexBio) or 50 μM dimethyl sulfoxide (DMSO) (Sigma‒Aldrich) and cultured for 48 hrs. This medium was replaced with normal medium, and the cells were allowed to culture for 72-96 hrs before the medium was replaced with serum-free medium containing only 1% penicillin‒streptomycin and allowed to culture for 24 hrs. These media were collected as conditioned media (CM) and used in subsequent experiments as either bleomycin-conditioned media (BCM) or control (DMSO)-conditioned media (CCM).

### Conditional Knockdown with siRNA

MIN6 cells were seeded in 6-well plates (300,000 cells/well) and cultured for 24 hrs. The siRNA transfection mixture was prepared using Opti-MEM I Reduced Serum Media (Gibco, Billings, MT; Lot:1747285), Lipofectamine RNAiMAX reagent (Invitrogen Waltham, MA; Lot:2538674) and p21^Cip1^ (Santa Cruz Biotechnology Inc., Dallas, TX; sc:29428) at 50 nM according to the manufacturer’s instructions and incubated for 5 minutes at room temperature. Following incubation, fresh medium was added to the cells, the mixture was applied dropwise to each well, and a homogenous distribution was achieved. After 48 hrs, the cells were collected and the pellets were analyzed for gene expression by qPCR.

Alternatively, isolated islets from retired breeder *C57/BL6* mice were dispersed into individual pancreatic cells and cultured overnight in RPMI culture medium. After the overnight incubation, islets underwent siRNA transfection using Lipofectamine RNAiMAX and Opti-MEM for 48 hrs. Each well received a master mix containing either scrambled siRNA (control) or p21 siRNA. For each well, a master mix containing 50 µL Opti-MEM and 1.25 µL Lipofectamine was combined with 3 µL of 10 µM siRNA for a final concentration of 50 nM. An additional 50 µL of Opti-MEM were added to each well. After the 48-hr incubation, the medium was replaced with RPMI, which was subsequently changed every two days. Cells were harvested 4, 6, 8, 10, and 12 days post-transfection. Prior to collection, glucose-stimulated insulin secretion (GSIS) assays were performed, with supernatants frozen at -80°C for later ELISA-based insulin quantification. Total RNA was extracted from the harvested cells, reverse transcribed into cDNA, and analyzed by quantitative PCR to determine gene expression levels.

### *In vitro* Recombinant Protein Treatment

Approximately 300,000 MIN-6 cells were plated into 24-well plates with 500 µL of Dulbecco’s modified Eagle’s medium supplemented with 15% FBS. The cells were incubated at 37°C with 5% CO_2_ for 24 hrs to allow attachment. The concentrations of the following recombinant proteins were compared to those detected in plasma by immunoassay (https://www.proteinatlas.org): DUSP3 (1.3 µg/L), GDF15 (1.3 µg/L), IDE (0.31 µg/L), LSAMP (150 µg/L), HSP90 (71 µg/L), and GSTP1 (69 µg/L). The recombinant proteins were diluted RPMI 1640 media without FBS supplemented with 1% bovine serum albumin (BSA). Details are provided in **Supplementary Table 2**. The cells were treated for 48 hrs, followed by incubation for 48 hrs in normal media. After treatment, the cells were subjected to qPCR.

### Glucose-Stimulated Insulin Secretion (GSIS)

Kreb-Ringer HEPES solution was made at the following concentrations: 137 mM NaCl, 4.8 mM KCl, 1.2 mM KH_2_PO_4_, 1.2 mM MgSO_4_.7H_2_O, 2.5 mM CaCl_2_.2H_2_O, 16 mM HEPES sodium salt, and 0.1% BSA; all reagents were obtained from Sigma‒Aldrich. The solution was aliquoted into two separate tubes of high (20.2 mM/l) and low (2.8 mM/l) glucose using a 45% glucose solution (Corning). Islets and/or FACS-sorted cells from P21-tdTOM mice were preincubated at 5% CO_2_ at 37°C for 1 hr in low-glucose Krebs-Ringer solution. Low-glucose incubation was performed for 1 hr, and the supernatant was collected, followed by high-glucose incubation for another hr. Cells were collected at the end for normalization by RNA or DNA total content.

### Quantitative Real-Time PCR

RNA extraction was performed using a RNeasy Micro Kit (Qiagen, Cat No. 74004). For reverse transcription from RNA to cDNA, we used SuperScript™ IV Reverse Transcriptase (Invitrogen). The sequences of primers used are listed in **Suppl. Table 4**. Gene expression levels were measured using PowerUp™ SYBR™ Green Master Mix (Applied Biosystems) and dCT values for β-actin were calculated.

### Immunostaining, Image Acquisition, and Morphometric Analysis

The cells were fixed with 10% formalin and permeabilized with 0.3% Triton X-100 (Alfa Aesar). The cells were blocked with normal donkey serum at a 1:50 dilution in PBS. The sections were incubated overnight at 4°C with anti-Ki67 antibody. The next day, they were incubated with anti-insulin and then with fluorochrome-conjugated secondary antibodies and mounted with Fluoroshield™ with DAPI (Sigma Aldrich).

The antibodies used are listed in **Suppl. Table 5**. For each stain, all images were taken with the same settings in confocal mode using a Zeiss LSM 710 microscope such that comparisons across conditions could be made.

The intensity for proliferation analysis and the area for cell density were quantified using ImageJ.

Quantification of fluorescence intensity was performed through FIJI. TdTOMATO fluorescence was quantified, followed by classification into higher or lower signalling categories based on the mean integrated density. Lower signalling corresponded to values in the lower half of the distribution, whereas higher signalling represented values in the upper half. To address islet heterogeneity, immunostaining was adjusted using the corresponding tdTomato integrated density values. This method facilitated the comparison of the impact of increasing tdTOMATO intensity and, consequently, P21^CIP1^ expression and other marker β-cell markers: INSULIN, MAFA, NKX6, and PDX1.

Staining details are provided in the **Supplementary data**.

### Bulk RNA-Seq

Islet isolation was performed according to the protocol described by Gotoh, M., *et al*. RNA was extracted from isolated islets using a RNeasy Micro Kit and sent for sequencing to DNA Link (Los Angeles, CA). Data analysis was performed by the Bioinformatics and Biostatistics Core at Joslin Diabetes Center.

### Single-Cell RNA Sequencing (scRNA-Seq): Mouse

Our previously published [6] scRNA-Seq database (GSE149984) was reanalyzed with β-cells clustered into subpopulations using the expression of *Cdkn1a* and *Cdkn2a*. Briefly, ALZET mini-pumps with the insulin receptor antagonist S961 or PBS were surgically inserted into mice for 2 weeks as described by Aguayo-Mazzucato *et al*.^4^. Three groups of mice (n=4 animals per group) were used: control (PBS pumps), S961 (20 nmol/L/week for 2 weeks), and recovered (mice with S961 removed for 2 weeks) groups. For single-cell RNA sequencing (scRNA-seq), islets were isolated from all mice in each group on the same day, cultured overnight, and dispersed. Single-cell transcriptomic analysis was performed using the 10X Genomics Chromium Single-Cell Gene Expression Assay Core at Brigham and Women’s Hospital. Gene UMI counts were generated by cellranger 3.0.2. Differential gene expression was performed by using the R packages, scatter and scran. *Cdkn1a-* and *Cdkn2a*-positive and Cdkn2a-negative cells from all three conditions were defined by using density estimates for each gene using a Gaussian finite mixture model from the R package Mclust [33]. *Ins2* expression was used to identify β-cell clusters. Differential gene expression was assessed using linear modelling with limma [34]. A moderated F test was performed to detect genes that were differentially expressed between any two of the 2 groups (*i.e*. *Cdkn1a*-only positive, *Cdkn2a*-only positive and *Cdkn1a*/*Cdkn2a*-double negative). Cell clustering was performed by using the R package Seurat. We used the QC metrics to filter cells that had fewer than 1000 UMIs (low-quality cells), fewer than 500 features (low-quality cells), and more than 10% mitochondrial UMIs (dying cells). Quality control (QC) after filtering revealed a median UMI of 6568 in the control group, 9826 in the S961 group, and 8415 in the S961 recovery group. The median number of genes per cell was as follows: 2019 genes in the control group, 2861 in S961, and 2409 in the recovered group. The percentage of mitochondrial DNA had a median of 5% in the control group, 3.5% in the S961 group, and 4.9% in the recovery group. There were 2939 β-cells in the control group, 2896 cells in the S961 group, and 2513 cells in the recovery group.

For βSenMayo, the list of genes described in [15] was assessed in different senescent subpopulations of β-cells from the control condition. Only those genes with quantifiable gene reads in all 3 subpopulations were included in the analysis to exclude those with extremely low expression in β-cells.

Reaction pathways were analysed using REACTOME (www.reactome.org), where all nonhuman genes were converted to their human equivalents for analysis.

A detailed description of the bioinformatic analysis of the scRNA-seq data can be found in the **Supplementary Methods**. The accession number for the scRNA-seq data is GSE149984.

### scRNASeq of Human Islets

Pancreatic islets from Prodo Lab were digested with StemPro™ Accutase™ Cell Dissociation Reagent at a concentration of 1 ml/1,000 islets for 10 minutes at 37°C. All dissociated cells were filtered through a 40 µm Flowmi® Cell Strainers (SIGMA) and counted using AO/PI (acridine orange/propidium iodide) Cell Viability Kit for Luna-FL automated cell counter. Cells were fixated prior to scRNA-seq according to 10X Genomics protocol CG000478. After determining cell concentration of the fixed sample by AO/PI Cell Viability Kit for Luna-FL automated cell counter pre-warmed Enhancer (10x Genomics PN-2000482) was added and samples stored for up to 1 week. For single cell library preparation and sequencing the cell viability was assessed on a LUNA FX7 automated cell counter (Logos Biosystems), and up to 12,000 cells from each suspension were loaded onto one lane of a 10x Genomic Chip [G]. Single cell capture, barcoding, and library preparation were performed using the 10x Genomics Chromium X platform [CITE https://www.nature.com/articles/ncomms14049] version 3.1 NEXTGEM chemistry and according to the manufacturer’s protocol (#CG000315). cDNA and libraries were checked for quality by Tapestation 4200 (Agilent) and Qubit Fluorometer (ThermoFisher), quantified by KAPA qPCR, and sequenced on an Illumina NovaSeq X+ 10B 100 cycle flow cell lane, with a 28-10-10-90 asymmetric read configuration, targeting 6,000 barcoded cells with an average sequencing depth of 50,000 reads per cell. Illumina base call files for all libraries were converted to FASTQs using bcl2fastq v2.20.0.422 (Illumina) and FASTQ files associated with the gene expression libraries were aligned to the GRCm38 reference assembly with vM23 annotations from GENCODE (10x Genomics mm10 reference 2020-A) using the version 8.0.0 Cell Ranger multi pipeline (10x Genomics).

### FFPE Block DV200 Evaluation

Prior to the spatial transcriptomics assay, the FFPE blocks were assessed for RNA quality by DV200 scores. Two to four 5µm FFPE sections were collected in RNAse free conditions and RNA was extracted using Qiagen’s RNeasy Micro Kit (Cat. No. / ID: 74004). RNA quality was then quantified using a Tapestation High Sensitivity RNA ScreenTape (Agilent). DV200 scores were calculated as the percentage of the RNA sample above 200 bases in length, as determined using Agilent TapeStation Analysis software. DV200 score of more than 50% was achieved for all FFPE blocks.

### CODEX/ PhenoCycler Slide Preparation, Setup, and Data Acquisition

The slides were baked at 60°C for 30 minutes. After baking, slides were immersed in xylene twice for 5 minutes each, followed by incremental rehydration in 100%, 95%, 70%, and 50% ethanol for 3 minutes each. The slides were then immersed in water for 3 minutes and moved to 1x antigen retrieval buffer (pH 9.0) (AR9, Akoya Biosciences), for antigen retrieval at 100 °C for 15 minutes using the TintoRetriever (BioSB).

Following antigen retrieval, the slides were cooled in the retrieval buffer to RT and washed in nuclease-free water for 5 minutes. Slides were then processed according to the PhenoCycler-Fusion User Guide (PD-000011 REV M, Akoya Biosciences), starting from step 4 on page 49. Briefly, the slides were washed in Hydration Buffer, equilibrated in Staining Buffer, and incubated overnight at 4°C with a 38-marker antibody cocktail prepared in Blocking Buffer. The slides were subsequently washed in Staining Buffer, gently fixed with Post-Staining Fixing Solution, washed in 1x PBS, and incubated in ice-cold methanol for 5 minutes. The slides were then washed in 1x PBS, fixed with Final Fixative Solution for 20 minutes, washed three times in 1x PBS, and immersed in Storage Buffer prior to the PhenoCycler Fusion run.

The experimental protocol was set up using the PhenoCycler Experiment Designer (Version 2.1.0, Akoya Biosciences). A reporter plate containing fluorescently labelled barcode reporters, as per the experimental design, was prepared following instructions on page 73 of the PhenoCycler-Fusion User Guide (PD-000011 REV M, Akoya Biosciences). Slides were prepared for the PhenoCycler-Fusion instrument according to the steps outlined in the PhenoImager-Fusion User Guide (PD-000001 Rev N, Akoya Biosciences). Briefly, the slide was moved from Storage Buffer to 1x PBS and incubated for 10 minutes. After incubation, a Flow Cell (Akoya Biosciences) was attached to the sample slide using the Flow Cell Assembly Device (Akoya Biosciences). The slide with the attached Flow Cell (Sample Flow Cell) was then placed in 1x PhenoCycler buffer for 10 minutes. PhenoCycler Fusion software (Version 2.2.0) was used to set up the imaging run on the PhenoCycler-Fusion, following the steps on page 57 of the PhenoImager-Fusion User Guide (PD-000001 Rev N, Akoya Biosciences). Reagents were prepared and loaded into the appropriate reagent reservoirs on the instrument, and the pre-prepared reporter plate was loaded into the PhenoCycler. A new PhenoCycler run was initiated using the experimental protocol design. A blank flow cell was loaded into the Flow Cell Slide Carrier, and all software prompts during the pre-flight routine were followed. The Sample Flow Cell was then loaded into the carrier and a leak check was performed. Scan regions were selected following automated sample finding and imaging was started. Upon completion, the Sample Flow Cell was placed in Storage Buffer at 4°C. The generated QPTIFF data file was used for downstream image analysis.

### Xenium

Genes for Xenium panels were selected based on literature review and initial scRNA-seq and Visium analysis. Given a list of gene targets ordered by level of importance, a custom 300 gene panel was designed using the 10x Genomics Xenium Panel Designer web-based tool (cloud.10xgenomics.com,) using Visium CytAssist and Fixed scRNA-seq (10x Genomics) data generated from matched samples as a reference to gauge anticipated transcript abundance and ubiquity. Due to the highly abundant genes targeted, such as INS, GCG, and SPINK2, the panel design process was run in an iterative fashion where (a) each target initially excluded due to high expression was reassigned 1 probe, then (b) the designer was run to generate probeset assignments for all other genes to accommodate the high-expressers. This process was repeated until the panel designer no longer excluded desired highly expressed genes.

FFPE Tissue sections of 5µm thickness were placed on the Xenium slide (10x Genomics, Cat. No. 1000660) such that 2 sections filled the 10.45 x 22.45mm capture area. The tissue was deparaffinized, permeabilized, de-crosslinked, and stained with probes according to manufacturer specifications (protocol CG000580, user guide CG000582). Slides were loaded onto the Xenium Analyzer and *in situ* gene expression data were generated according to the manufacturer’s user guide (CG000584, RevG). After the Xenium run, data were exported from the analyzer for analysis, and Xenium slides were subject to H&E staining according to the manufacturer demonstrated protocol (CG000613) and imaging via Nanozoomer SQ (Hamatsu). Following Xenium, multiplexed antibody-based immunofluorescence staining and imaging was performed using the Phenocycler-Fusion (Akoya Biosciences) according to a custom protocol (https://dx.doi.org/10.17504/protocols.io.q26g71rwqgwz/v1)].

### β-Gal Activity Assay

To measure senescence-associated β-galactosidase (SA-βGal) activity, we used the *Dojindo Cell Count Normalization Kit (C544)* combined with the *Cellular Senescence Plate Assay Kit – Spider-βGal (SG05)* following the manufacturer’s protocol for combined analysis.

### Flow Cytometry 1. β-Gal^-^ and β-Gal^+^ Human and Mouse Cells

Human and mouse β-cells were sorted using flow cytometry based on β-Gal activity to divide them into senescent (β-Gal^+^) and nonsenescent (β-Gal^-^) populations. Β-Gal activity was measured using an Enzo cellular senescence live-cell senescence assay kit (ENZ-KIT130-0010) following the manufacturer’s instructions while optimizing the substrate incubation time to 1 hr. Sorting was performed using a DakoCytomation MoFlo Cytometer or FACSAria in the Joslin DRC Flow Cytometry Core. Primary islets were incubated with antibodies against CD45 and CD11b to eliminate immune cells. <u>**2. p21-tdTom mice**</u>. Islets from *p21^Cip1^-*tdTomato mice were isolated, dispersed into single cells, and sorted using a MoFlo Cytometer based on Tomato Red fluorescence. The cells were collected in media, plated in 96-well plates overnight, and assessed for glucose-stimulated insulin secretion the following day. The results were normalized to the DNA quantity. A detailed protocol for staining for *p16^Ink4+^* is provided in the **Supplementary Methods**. The gating criteria are shown in **Suppl. Fig. 8**, and 90% enrichment of β-cells was achieved, as previously published [3].

### Proteomics of Human Conditioned Media From β-Gal-/β-Gal^+^ Human Cells and Control and JAK1/2i-Treated Islet Cells

After sorting, β-Gal^+^ and β-Gal^-^ cells were plated, and conditioned media were generated to compare the SASP secreted by either cell type. Conditioned media from control and pharmacologically treated cells were generated as described above. The generated conditioned media were analysed using SomaScan at the Genomics Proteomics Core (BIDMC).

The Somascan data were normalized for hybridization, and to obtain the same median, the data were then log_2_ transformed. PCA was performed to assess differences among samples. Associations between the experimental variables and the top 1-6 principal components were tested. PCA plots were adjusted using the empirical Bayes method [35]. Limma was used to identify differentially secreted proteins, and moderated t tests were used to detect significant differences by paired analysis.

Human proteomic βGal data are available in the GSE150285 dataset, and human data for the control, JAK1/2i datasets are available in the GSE162521 dataset.

### Quantification and Statistical Analysis

The data are shown as the mean ± SEM, and p values < 0.05 were considered significant. For statistical analysis, unpaired Student’s t tests and 2-way ANOVA were used to compare groups. Normality and log normality analyses were performed, and nonparametric statistics (Kruskal‒Wallis test, Mann‒Whitney test, and Wilcoxon test) were performed when samples did not meet the criteria for a normal distribution. Prism software was used for graphs and statistical analysis (significance and distribution). Data outliers were determined using the Grubbs outlier test or a deviation of more than 2 SDs from the mean.

## Acknowledgements

The authors thank Jonathan Dreyfuss from Joslin’s Bioinformatic Core for assisting with the data analysis; Angela Wood and Alison Marotta from the Flow Cytometry Core and Erin Keating of Animal Facilities; Stephan Kissler and Taylor Roberts from the JDC Mouse Genomic Core; and Susan Bonner-Weir for insightful discussion, critical reading of the manuscript and support in obtaining human islets from the IIDP.

This work includes data and/or analyses from **HumanIslets.com** funded by the Canadian Institutes of Health Research, JDRF Canada, and Diabetes Canada (5-SRA-2021-1149-S-B/TG 179092) with data from islets isolated by the Alberta Diabetes Institute IsletCore with the support of the Human Organ Procurement and Exchange program, Trillium Gift of Life Network, BC Transplant, Quebec Transplant, and other Canadian organ procurement organizations with written informed donor consent as approved by the Human Research Ethics Board at the University of Alberta (Pro00013094).

## Funding

This study was supported by Institutional Startup Funds to CAM, National Institutes of Health (NIH) grants 1R01DK132535 to CAM, P30 DK036836 to Joslin Diabetes Center (Cores), Thomas J Beatson Jr Foundation grant 2020-010, and the Richard and Susan Smith Family Foundation Award to CAM. SP was funded by Award Number 1R25DK113652 from the NIH NIDDK. SL was funded by R25 DK140752 - Joslin-BIDMC Post-Bac Program. JLK and TT were funded by NIH grant R37 AG13925, Hevolution grant HF-GRO-23-1199148-3, and the Connor Fund, Robert and Theresa Ryan, and the Noaber Foundation. KI was supported by The Mampei Suzuki Diabetes Foundation and a Mary K. Iacocca Fellowship. This work received support from the NIH/NIA U54AG075941 to the SenNet Initiative, KAPPSen Tissue Mapping Center. The content is solely the responsibility of the authors and does not necessarily represent the official views of the National Institute of Diabetes and Digestive and Kidney Diseases, the National Institute on Aging, or the National Institutes of Health.

Human islets were provided by the NIDDK-funded Integrated Islet Distribution Program (IIDP) at the City of Hope (NIH grant no. 2UC4DK098085) and from Prodo Laboratories, Inc., in Aliso Viejo, CA.

## Author contributions

CA, PC, KI, FH, SP, and SS acquired, analyzed, and interpreted the data and wrote, edited, and revised the manuscript. AM generated the *p21^Cip1^-*tdTomato mice with the assistance of the mouse genomic core at Joslin, and HP analyzed the proteomic, scRNA-seq, and RNA-seq data. CAM designed the project; acquired, analyzed, and interpreted the data; and wrote, edited, and revised the manuscript.

SenNet KAPPSen members: JAD, DB, SD, SE, AP, FGC, JC, GAK, NM, PR, TT, JLK, VG, JLW contributed to organ retrieval, scRNASeq, CODEX, and Xenium on whole human pancreas and human islets.

All authors read, revised and approved the manuscript.

## Competing interests

Part of the work in this manuscript has been submitted for a patent JDP-216 on January 16, 2024, as a US Provisional, 63,621,239.

