## Supplementary data for "A human and mouse subpopulation of senescent β-cells induces pathologic dysfunction through targetable paracrine signaling"

SUPPLEMENTARY DATA TO Iwasaki K, Carapeto P. et al.

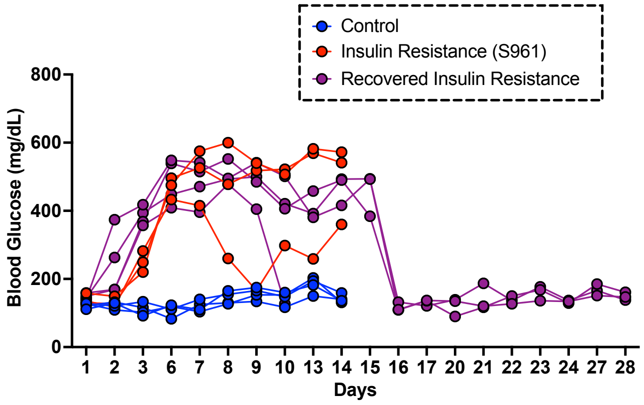


**Suppl. Fig. 1 related to Fig. 1A-E** Blood glucose levels in mice treated with S961 with and without 2 week recovery period. These data have been previously published in (6)

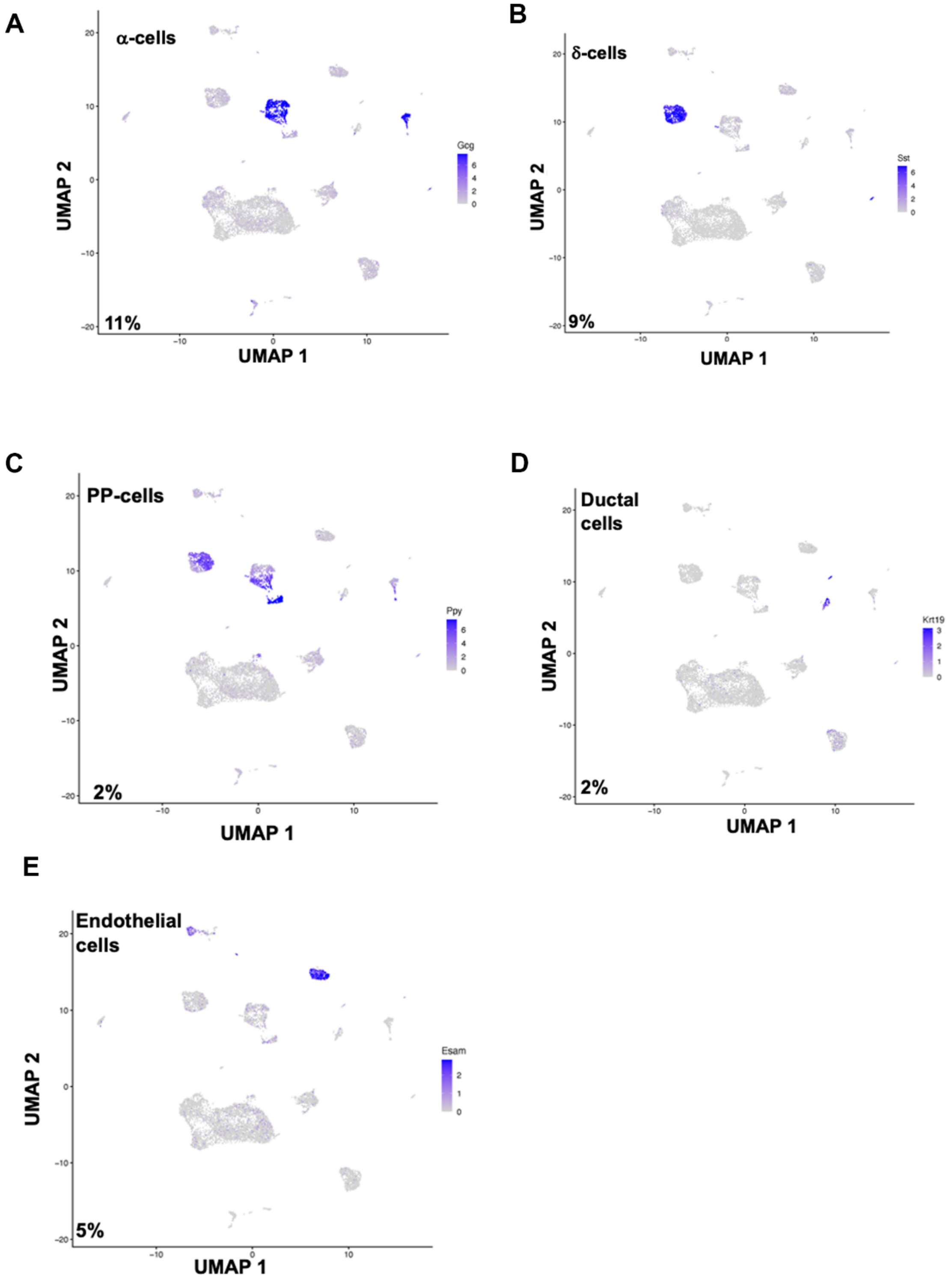


**Figure S2 related to figure 1B:** UMAP of non-β-cell subpopulations after scRNASeq as determined by signature genes in mice. (**A**) Alpha cells by *Gcg* expression; (**B**) Delta cells by *Sst* expression; (**C**) PP cells by *Ppy* expression; (**D**) ductal cells by *Krt19* expression and (**E**) endothelial cells by *Esam* expression.

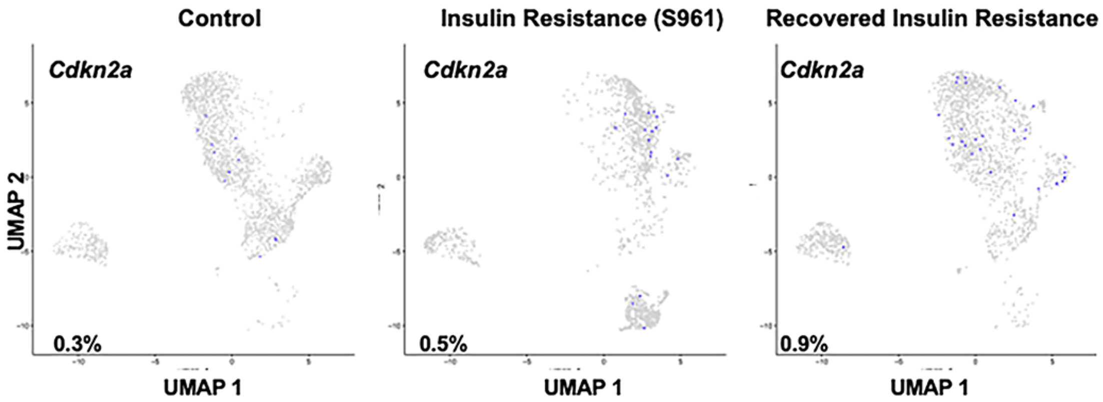


**Fig S3 related to Fig. 1C.** UMAP showing the percentage of *Cdkn2a*$^{+}$ β-cells under three different metabolic conditions as determined by scRNASeq in 8-9 month old C57Bl/6 mice.

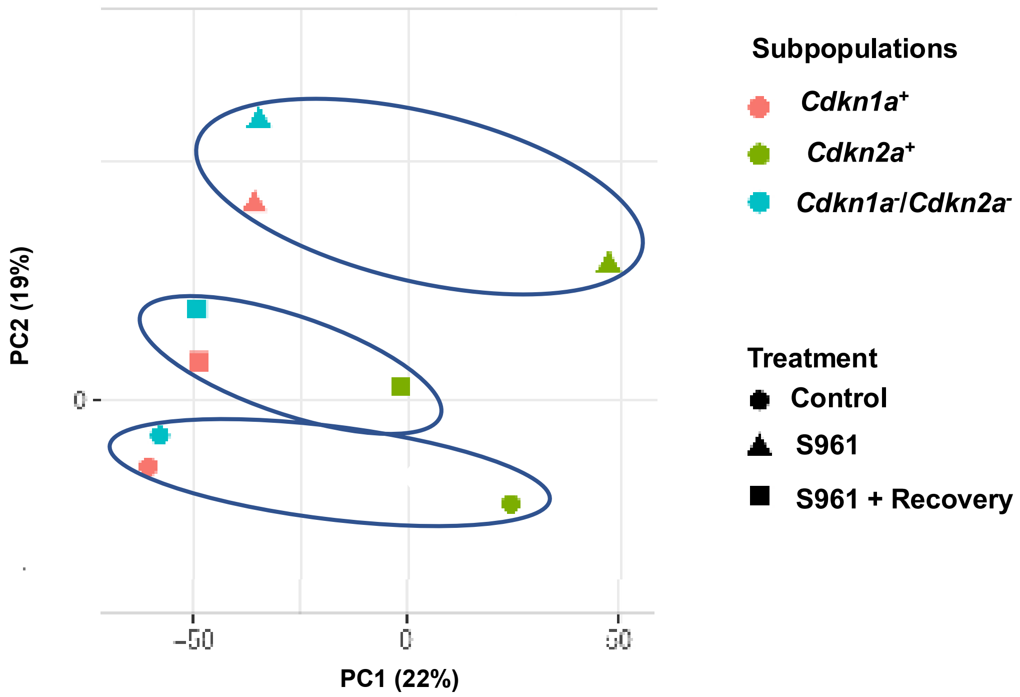


**Fig S4 related to Figs. 1D, E.** PCA analysis of the z-scores of mouse β-cells shown in Figs. 1D, E, shows significant clustering by treatment, Therefore, results are shown for all three treatments in the 3 cell subpopulations.

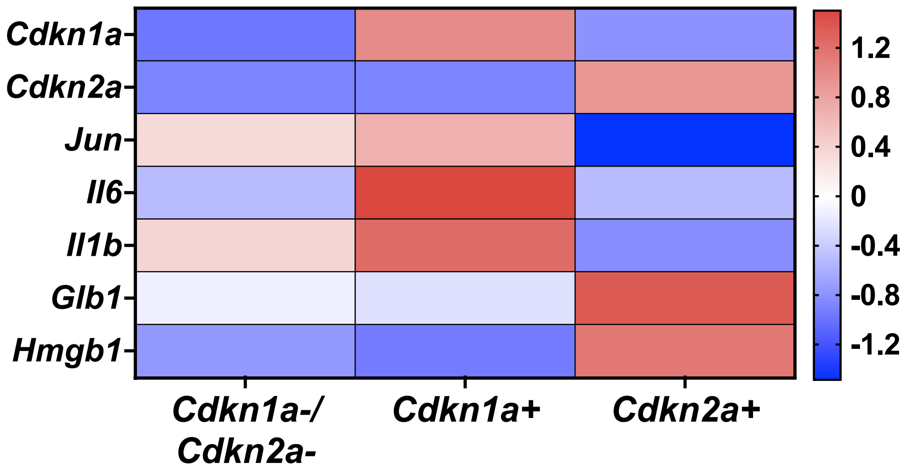


**Fig S5 related to Fig 1.** Heatmap of senescence marker genes.

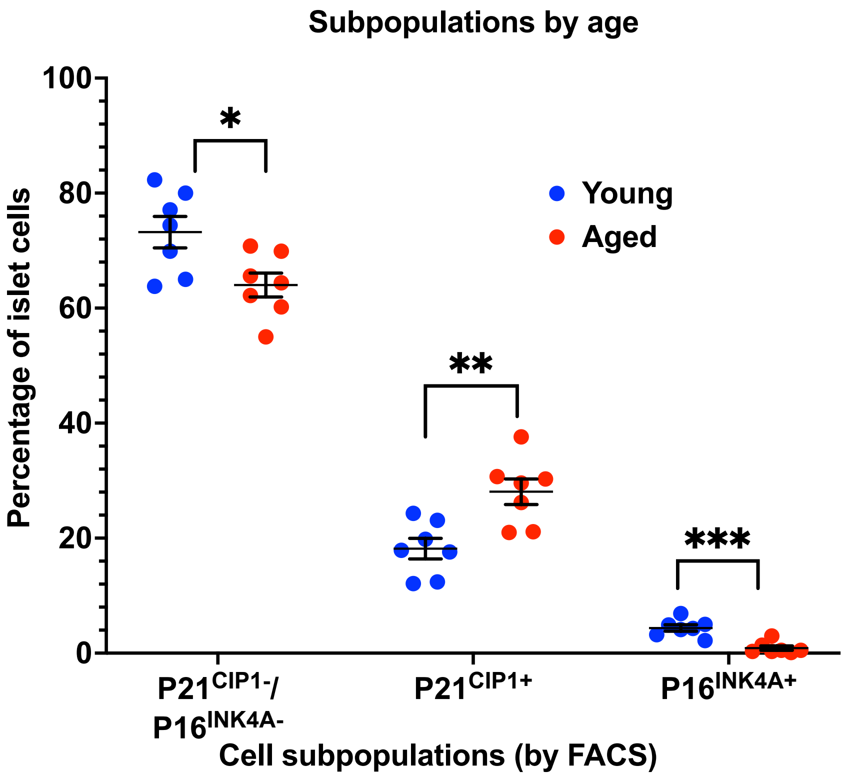


**Fig. S6 related to Fig. 2**. Percentage of *P21CIP1*$^{+}$ (encoded by *Cdkn1a*), *P16INK4A*$^{+}$ (encoded by *Cdkn2a*) and double *P21CIP1*$^{-}$/ *P16INK4*$^{-}$ islet cells by flow cytometry from islets isolated from young (3 months) and aged (18 months) *p21*$^{Cip}$-tdTomato mice and stained for *p16*$^{Ink4a}$, n=4 independent experiments

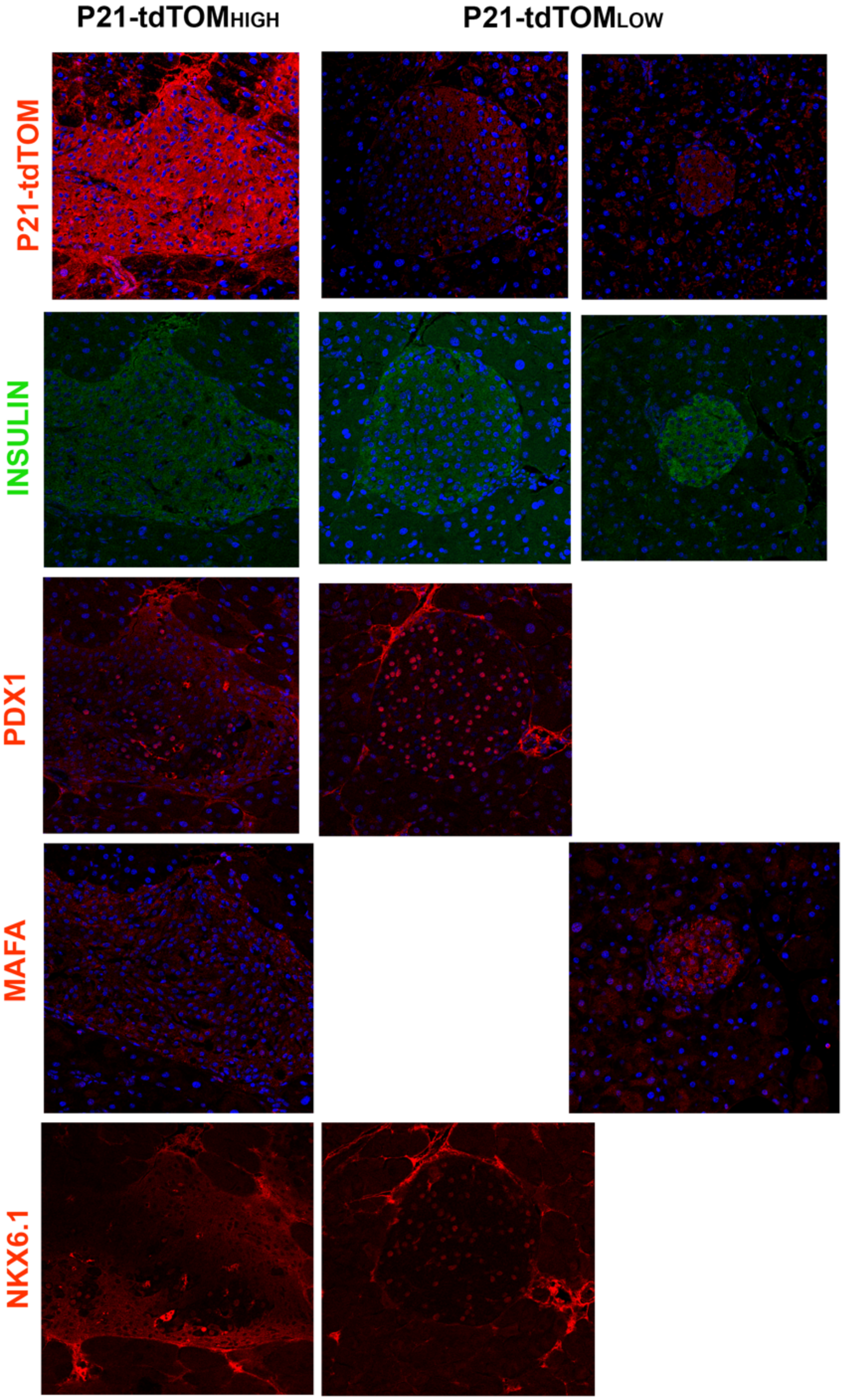


**Figure S7 related to figure 2G.** Loss of β-cell identity at the protein level. Quantification of protein intensity in $tdTOM^{HIGH}$ islets normalized to $tdTOM^{LOW}$. Each point represents an individual islet from two P21td-TOM positive mice.

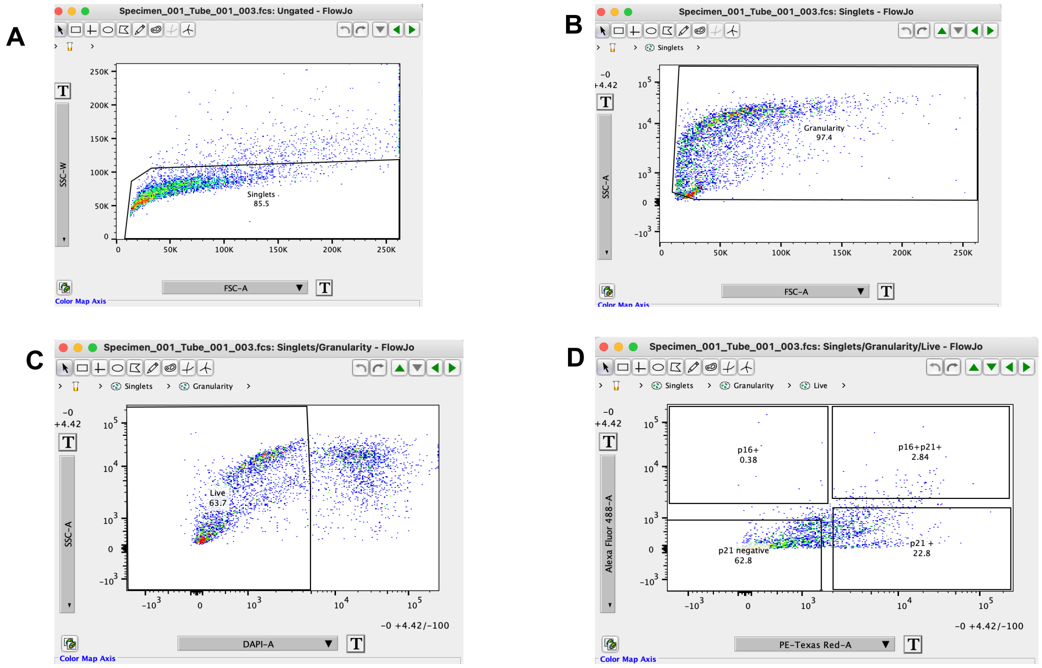


**Suppl. Fig. 8 related to Fig. 4K and 5K**. Flow cytometry gating strategy for identification of senescent subpopulations. A. Gating to identify single cells followed by B. Gating based on granularity, C. Gating to identify live cells through the exclusion of DAPI and D. Identification of 3 cell subpopulations based on the intensity of dTomRed for P21CIP1 and Alexa fluor-488 for P16INK4A.

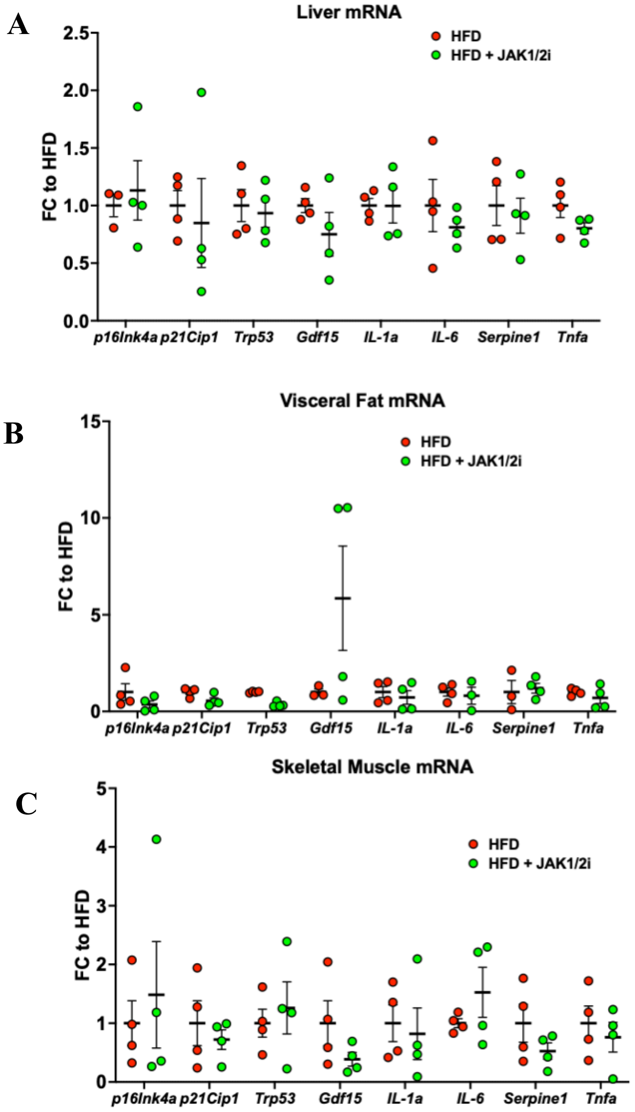


**Suppl. Fig. 9 related to Fig. 5. Differential gene expression of senescence and SASP markers across peripheral tissues can be partially recovered by senotherapeutics.** Liver, visceral adipose tissue from the peritoneum, and skeletal muscle from the gastrocnemius were isolated from mice at the end of treatment with n=3 for Control, n=4 for high-fat diet (HFD), and n=4 for HFD+JAK1/2i. All samples were assayed by qPCR for senescence genes p16Ink4a, p21Cip1, and Trp53 and SASP genes Gdf15, Il-1a, Il-6, Serpine1, and Tnfa. All bars represent mean fold change ± SEM A) qPCR of isolated liver from HFD, and HFD+JAK1/2i mice with fold-change to HFD; *$p<0.05$ and **$p<0.01$ by unpaired t-test relative to HFD. B) qPCR of isolated visceral adipose tissue from HFD, and HFD+ JAK1/2i mice with fold-change to HFD; **$p<0.01$ and ***$p<0.001$ by unpaired t-test relative to HFD. C) qPCR of skeletal muscle from HFD, and HFD+ JAK1/2i mice with fold-change to HFD; **$p<0.01$ by unpaired t-test relative to HFD.

**Supplementary Table 1.** Percentages of senescent cell subpopulations in different islet cell types in control, insulin resistance (S961) and recovered insulin resistance (S961R) groups.

| Group | Cdkn1a$^-$/Cdkn2a$^-$ | Cdkn1a$^+$/Cdkn2a$^-$ | Cdkn1a$^-$/Cdkn2a$^+$ |
|---|---|---|---|
| **Alpha cells** | | | |
| Control | 86.7 | 13.3 | 0.0 |
| S961 | 73.7 | 25.7 | 0.3 |
| S961+R | 82.9 | 16.4 | 0.7 |
| **Delta cells** | | | |
| Control | 78.2 | 21.8 | 0.0 |
| S961 | 62.7 | 37.3 | 0.0 |
| S961+R | 75.6 | 24.4 | 0.0 |
| **PP cells** | | | |
| Control | 75.0 | 25.0 | 0.0 |
| S961 | 67.7 | 31.3 | 1.0 |
| S961+R | 72.5 | 27.5 | 0.0 |

**Supplementary Table 2. Selected SASP recombinant proteins for in vitro treatment of pancreatic islets**

| Protein | Human/Mouse | Initial Dry Mass | Source | Catalog number |
|---|---|---|---|---|
| GDF-15 | Mouse | 25 ug | R&D Systems | 8944-GD-025 |
| IDE | Mouse | 20 ug @0.07ug/ul | OriGene | TP525567 |
| LSAMP | Mouse | 20 ug @0.15ug/ul | OriGene | TP523437 |
| DUSP3 | Mouse | 20 ug @0.05ug/ul | OriGene | TP501720 |

**Supplementary Table 3. Human Islets Donor Information**

| Age | Gender | T2D | BMI | Use |
|---|---|---|---|---|
| 27 | M | No | 25.3 | GSIS, qPCR senolytic/senomorphic, proteomics, staining |
| 26 | M | No | 24.2 | GSIS, qPCR senolytic/senomorphic, proteomics, staining |
| 21 | M | No | 27.17 | GSIS, qPCR senolytic/senomorphic, proteomics, staining |
| 22 | M | No | 31.9 | GSIS, qPCR senolytic/senomorphic, proteomics, staining |
| 24 | M | No | 33.6 | GSIS, qPCR senolytic/senomorphic, proteomics, staining |
| 42 | M | No | 30.3 | qPCR senolytic/senomorphic |
| 29 | M | No | 26.2 | qPCR senolytic/senomorphic, GSIS |
| 64 | M | No | 29.1 | qPCR senolytic/senomorphic, GSIS |
| 34 | | No | 24.6 | qPCR senolytic/senomorphic |
| 19 | F | No | 22.3 | qPCR senolytic/senomorphic, GSIS |

| | | | | |
|---|---|---|---|---|
| 58 | M | Yes | 38.3 | Perifusion |
| 49 | F | Yes | 42.2 | qPCR senolytic/senomorphic, GSIS |
| 55 | F | Yes | 42.2 | qPCR senolytic/senomorphic, GSIS |
| 60 | F | Yes | 29.9 | qPCR senolytic/senomorphic |
| 21 | M | No | 28 | Xenium, CODEX |
| 71 | F | No | 26 | Xenium, CODEX |
| 32 | M | No | 28 | Xenium, CODEX |
| 54 | M | No | 23.9 | Xenium, CODEX |
| 28 | F | No | 17.7 | Xenium, CODEX |
| 22 | M | No | 22.4 | Xenium, CODEX |
| 23 | F | No | 17.2 | Xenium, CODEX |
| 40 | F | No | 22.97 | Xenium, CODEX |
| 37 | F | No | 23.5 | Xenium, CODEX |
| 47 | M | No | 21.8 | Xenium, CODEX |
| 69 | F | No | 18 | Xenium, CODEX |
| 35 | M | No | 23.4 | Xenium, CODEX |
| 39 | M | No | 24.7 | Xenium, CODEX |
| 43 | M | No | 14.4 | Xenium, CODEX |
| 62 | F | No | 23 | scRNA-seq |
| 46 | M | No | 23.1 | scRNA-seq |
| 42 | F | No | 29.3 | scRNA-seq |
| 50 | F | No | 28.3 | scRNA-seq |
| 57 | M | No | 30.9 | scRNA-seq |
| 59 | M | No | 24.5 | scRNA-seq |
| 58 | F | No | 31 | scRNA-seq |
| 64 | M | No | 25.5 | scRNA-seq |
| 47 | M | No | 28.3 | scRNA-seq |
| 57 | F | No | 30.2 | scRNA-seq |
| 58 | M | No | 28.9 | scRNA-seq |
| 34 | M | No | 24.6 | scRNA-seq |
| 61 | M | No | 29.5 | scRNA-seq |

**Supplementary Table 4. Primer Sequences**

| Gene | | Forward | Reverse |
|---|---|---|---|
| *β-actin* | Mouse | ACCGTGAAAAGATGACCCAG | GTACGACCAGAGGCATACAG |
| *p21Cip1* | Mouse | GCAGATCCACAGCGATATCC | CAACTGCTCACTGTCCACGG |
| *p16Ink4a* | Mouse | CCCAACGCCCCGAACT | GCAGAAGAGCTGCTACGTGAA |
| *Lsamp* | Mouse | CCCAAGACCTCCCAAGTTTAC | CCAGGTGATAACAGGTTCAGG |
| *Igf1r* | Mouse | ATTCTGATGTCTGGTCCTTCG | AGCATATCAGGGCAGTTGTC |
| *Il1a* | Mouse | TCAGCACCTTACACCTACC | GAGATAGTGTTTGTCCACATCC |
| *Il6* | Mouse | CAGAGGATACCACTCCCAAC | CAATCAGAATTGCCATTGCAC |
| *Insulin* | Mouse | CCTGTTGGTGCACTTCCTA | TCTGAAGGTCCCCGGGGCT |

| | | | |
|---|---|---|---|
| *Mafa* | Mouse | CTTCAGCAAGGAGGAGGTCATC | GCGTAGCCGCGGTTCTT |
| *Nkx6.1* | Mouse | GAAGAGAAAACACACCAGACC | CCGTCATCCCCAGAGAATAG |
| *Gck* | Mouse | TGAAACACAAGAAACTACCCC | CCACATCCATCTCAAAGTCC |
| *TBP* | Human | TGTGCACAGGAGCCAAGAGT | ATTTTCTTGCTGCCAGTCTGG |
| *P21CIP1* | Human | TGTCACTGTCTTGTACCCTTG | GCGTTTGGAGTGGTAGAAATC |
| *P16INK4A* | Human | CTTCGGCTGACTGGCTG | GCCTCCGACCGTAACTATTC |
| *IL6* | Human | AAAAGTCCTGATCCAGTTCCTG | TGAGTTGTCATGTCCTGCAG |
| *CCL4* | Human | TCCTCGCAACTTTGTGGTAG | TCCAGGTCATACACGTACTCC |

**Supplementary Table 5. Antibodies for Immunostaining**

| Antibody | Source | Catalog Number | Concentration |
|---|---|---|---|
| Anti-Insulin | Abcam | Ab195956 | 1:200 |
| Anti-HMGB1 | BioVision | A1005-50 | 1:400 |
| Anti-Ki67 | Abcam | Ab15580 | 1:500 |
| Anti-PDX1 | DSHB Antibodies | F6A11 | 1:50 |
| Anti-Nkx6.1 | DSHB Antibodies | F55A12 | 1:200 |
| Anti-MafA | Abcam | Ab26405 | 1:200 |
| Anti-tdTomato | Antibodies.com | A121690 | 1:250 |
| 488 Donkey anti-Guinea Pig IgG | Jackson ImmunoResearch Laboratories | 706-545-148 | 1:200 |
| 594 Donkey anti-Rabbit IgG | Jackson ImmunoResearch Laboratories | 711-585-152 | 1:200 |
| 594 Donkey anti-Goat IgG | Jackson ImmunoResearch Laboratories | 705-005-003 | 1:200 |

### Supplementary methods

#### scRNASeq Analysis

Gene UMI counts in droplets were generated by aligning reads to the mouse genome (mm10). To distinguish between droplets containing cells RNA and ambient RNA, we used Monte Carlo simulations to compute *p*-values for the multinomial sampling transcripts from the ambient pool (Lun et al. 2019). First, we assumed that some barcodes correspond to empty droplets if their total UMI counts were at or below 250, 150, and 200 for CTRL, S961 and S961RECOVERED respectively. We then called cells at a false discovery rate (FDR) of 0.1%, meaning that no more than 0.1% of our called barcodes should be empty droplets on average. The number of Monte Carlo iterations determined the lower bound for the *p*-values (Phipson and Smyth 2010). There were no non-significant barcodes bounded by iterations, which indicated we didn't need to increase the number of iterations to obtain even lower *p*-values. To estimate maximum ambient contamination, we computed the maximum contribution of the ambient solution RNA to the expression profile for cell-containing droplets (Lun et al. 2019). Firstly, we estimated the composition of the ambient pool of RNA based on the barcodes with total UMI counts less than or equal to 250, 150, and 200 for each gene in CTRL, S961 and S961RECOVERED respectively. Secondly, we computed the mean ambient contribution for each gene by scaling the ambient pool by some factor. Thirdly, we computed a *p*-value for each gene based on the probability of observing an ambient count equal to or below that in cell-containing droplets based on Poisson distribution. Fourthly, we combined *p*-values across all

genes using Simes' method (Simes 1986). We performed this for a range of scaling factors and identified the largest factor that yields a combined *p*-value above threshold 0.1 so that the ambient proportions were the maximum estimations.
We transformed and normalized the data for each sample using the R package sctransform (Hafemeister and Satija 2019). We performed PCA on the transformed data. Only the top 3000 variable genes were used as input.
**Cell clustering**. We constructed a K Nearest Neighbor (KNN) network of the cells based on the Euclidean distance in the space defined by the top principal components (PCs) that were selected by Horn's parallel analysis (Horn 1965). We refined the weights of the connection between pairs of cells based on their shared overlap in their local neighborhoods i.e. Jaccard similarity (Jaccard 1912). To cluster the cells, we applied modularity optimization techniques, i.e. the Louvain algorithm (Blondel et al. 2008).
**Quality control**. We visualized QC metrics as violin plots (Ilicic et al. 2016), which included the number of UMI (nCount), number of genes (nFeature), and percentage of mitochondrial UMI (percent_mt). QC metrics are reported in Main Methods section.

**Preparation of islets from p21-tdTom mice for FACS of $p21^+$/$p16^+$ cell populations**
p21-tdTom mice were used for islet collection and isolation was done following protocol described in Gotoh, M., *et al*. Islets were dispersed by incubation with TrypLE™ Express for 15 minutes in a 37°C water bath with vortexing every 3 minutes and resuspended in 4mL of 2% Fetal Bovine Serum (FBS) in 1x PBS. The dispersed islet cells were blocked using 1mL of Normal Goat and Normal Rat Serum from Jackson ImmunoResearch Labs in a 1:10 ratio in 1X PBS for 1 hour at 4°C. During this time the primary antibodies for p16 (Anti-CDKN2A/p16INK4a antibody ab54210 from abcam) was mixed from stock into a 1:20 ratio in 1X PBS then mixed again into a 1:20 ratio in 2% FBS in PBS. Secondary antibodies were mixed into a 1:100 ratio in 2% FBS in 1X PBS. Since the $p21^+$ cells of the dTomato mice fluoresce at a red 581λ wavelength, the secondary antibody for $p16^+$ cell used was fluorescent for green at 488λ wavelength (AlexaFluor 488 Donkey anti-Rabbit). After blocking the solution was spun down at 1600RPM for 2 minutes and had the supernatant removed and replaced with 100uL of the primary antibody solution and left to incubate for 30 minutes at 4°C. Then, the solution was spun down at 1600RPM for 2 minutes and washed with 10mL of 2% FBS in 1X PBS. The solution was spun down again at 1600RPM for 2 minutes and the supernatant was removed and replaced with 300uL of secondary antibody solution and left to incubate for 15 minutes at 4°C. The solution was spun down at 1600RPM for 2 minutes and the dispersed islet cells were resuspended 2% FBS in 1X PBS for FACS sorting using DAPI as a viability stain. Unstained samples followed all washes and solution replacements using 2% FBS in 1X PBS only.

**Proteomics**
After sorting, βgal-positive and βgal-negative primary mouse β-cells were plated in 96-well plates and incubated in serum-free islet media to generate conditioned media which was analyzed using SomaScan at the Genomics Proteomics Core (BIDMC). Data analysis, including, was conducted using R. Graphs were generated using GraphPad Prism.

**Immunohistochemistry of mouse pancreas tissues**

Paraffin-embedded blocks were prepared from tdTom+ and C57BL/6 mice pancreases and the slides were cut at 4 microns. Deparafinization step was done in the following order: Xylene-14 min; 100 EtOH- 8 min; 95% EtOH-6 min; 75% EtOH-10 min and $ddH_2O$ 5 min. Slides were washed for 10 min in 2% Lamb serum containing PBS. Later avidin was applied for 30 min followed by PBS (2% Lamb) and biotin for 30 min. After another washing step with PBS (2%

Lamb) slides were incubated in Normal Donkey Serum (2% in PBS) for 30 minutes and then specific primary antibodies were incubated overnight at 4°C. On day 2 the slides were brought to RT for 1 hour then washed with PBS (2% Lamb) for 10 min. Afterwards respective secondary antibodies were applied for 1 hour at RT. Washing step with PBS followed then the second primary antibody was applied and incubated overnight at 4°C. On day 3 the slides were brought to RT for 1 hour then washed with PBS (2% Lamb) for 10 min. Afterwards respective secondary antibodies were applied for 1 hour at RT. Washing step with PBS followed then the sections were mounted with Fluoroshield with DAPI mounting media. Images were taken using confocal mode on a Zeiss LSM 710 microscope.
